# Spatial analysis reveals a novel inflammatory tumor transition state which promotes a macrophage-driven induction of sarcomatoid renal cell carcinoma

**DOI:** 10.1101/2025.06.12.656421

**Authors:** Allison M. May, Suguru Kadomoto, Claire Williams, Alex C. Soupir, Stephanie The, Jake J. McGue, Tyler Robinson, Greg Shelley, Mitchell T. Hayes, Brooke L. Fridley, Jodi A. Balasi, Paolo M. Ramos Echevarria, Jasreman Dhillon, Srinivas Nallandhighal, Satwik Acharyya, Liying Chen, Jessica Aldous, Nathan Schurman, Veera Baladandayuthapani, Timothy L. Frankel, Simpa S. Salami, Brandon J. Manley, Rohit Mehra, Aaron M Udager, Evan T. Keller

## Abstract

Sarcomatoid renal cell carcinoma (sRCC) is an aggressive transdifferentiation of epithelioid clear cell RCC (ccRCC) tumors that shows heightened response to immunotherapy. The underlying biology leading to sarcomatoid transformation and mechanisms contributing to immunotherapy response are not well understood. Novel single cell spatial techniques were used in ccRCC and sRCC tumors from 40 patients to understand the spatial sRCC transformation and corresponding immune changes. A transcriptional transition state in epithelioid ccRCC cells along a continuum to mesenchymal sRCC was identified which expresses high levels of pro-inflammatory cytokines and an immune infiltrate. *In vitro* studies demonstrated that M2-like macrophages, recruited to the tumor by the transition state, induce full transition to the sarcomatoid state. A combination of increased PD-L1 expression and T-cells recruited by the transition state was observed, consistent with the increased immunotherapy response. This study enriches our understanding of the mechanisms leading to development and immune responsiveness of sRCC paving the way for novel approaches to diminish RCC progression.

## INTRODUCTION

Renal cell carcinoma (RCC) is diagnosed in approximately 81,000 people in the United States annually(1) and while often curable when localized, in the advanced stage RCC has very poor prognosis (2, 3). Sarcomatoid RCC (sRCC) is one of the most aggressive forms of renal cancer, occurring in up to 20% of advanced renal tumors, and with median survival as low as 6 – 13 months (4). sRCC develops as a transformation of the primary tumor, most commonly clear cell RCC (ccRCC), and appears histologically as patches of pleomorphic and spindle-shaped cells, intermixed with patches of the epithelioid parental tumor cell type. Because of intra-tumoral heterogeneity in these tumors, it is believed that sRCC may be underdiagnosed (5).

sRCC is thought to develop through an epithelial to mesenchymal transition (EMT) (6, 7). EMT, a well described biologic pathway in which cells lose epithelial properties and gain mesenchymal properties, is often triggered by signaling from the microenvironment (8, 9). Prior studies exploring molecular factors associated with sarcomatoid transformation in RCC have identified genomic mutations in *TP53*, *BAP1,* or *NF2* amongst others (10, 11, 12, 13) may play a role, and others have speculated hypoxia or TGFβ signaling may induce EMT as both are increased in sRCC tumors (12, 13, 14). However, the initiating factors and dynamics of the sarcomatoid transformation predominantly remain unknown.

Despite the limited understanding of the molecular underpinnings of sRCC, recent trial data has unearthed an intriguing finding: sRCC tumors have increased sensitivity to immunotherapy compared to ccRCC without sarcomatoid changes (15, 16). This is a paradoxical finding since substantial evidence shows EMT signaling typically leads to an immunosuppressive state in cancer (8, 17, 18) which would not typically facilitate immunotherapy response that is dependent on cytotoxic T cell infiltrate (19). Understanding this paradox in sRCC is an increasingly important key to understanding immune response of RCC as a whole, as immunotherapy has become a mainstay of RCC management both in the metastatic and adjuvant setting (20, 21), however, only a subset of tumors show an objective and durable response and there are no biomarkers to predict this response (21).

The majority of prior molecular studies of sRCC have compared ccRCC and sRCC on a binary level; however, it is known that EMT processes occur on a continuum and recent evidence suggests there are hybrid EMT states along this continuum (22, 23). Based on these observations we hypothesized there is a spatial gradient of EMT from ccRCC through sRCC in RCCs and that these EMT states are associated with unique immune microenvironments. These spatially-identified immune alterations in the tumor microenvironment could provide clues to the mechanisms of transition from ccRCC to sRCC and the associated enhanced response to immunotherapy.

Herein, we used high resolution spatial biology techniques, incorporating transcriptomics and proteomics, to identify a unique hybrid EMT state within clear cell regions spatially near sarcomatoid regions of ccRCC/sRCC tumors which we believe represents a state of transition. We demonstrate features of this transition cell state can be found even in pure ccRCC tumors and are associated with poor prognosis, suggesting that this biology may drive progression in a broader subset of renal cancer. We identify a highly immune cell-infiltrated microenvironment in sarcomatoid and transition areas which appears to be driven by inflammatory features of the transitional hybrid EMT state. Finally, we show that transition and sarcomatoid areas within ccRCC tumors are particularly rich in M2-like macrophages and we demonstrate through *in vitro* experiments that crosstalk between RCC cells and SPP1 high macrophages drives the sRCC transformation and immune checkpoint upregulation.

## RESULTS

### Single cell spatial transcriptomic profiling of ccRCC/sRCC identifies a distinct tumor cell state in clear cells spatially abutting sarcomatoid regions

To examine the spatial landscape of sRCC, we identified 29 formalin-fixed paraffin-embedded (FFPE) whole block specimens from 20 patients with pathologically confirmed sRCC and selected blocks that contained both ccRCC and distinct sRCC elements (Figure 1A) for spatial transcriptomic profiling. This was performed at the single cell level on specimen 1 (Figure 1A-B) via the NanoString CosMx platform (24) using a 960 plex gene panel (full gene list in Table S1). Fields of view (FOVs) for spatial transcriptomics were selected based on histopathologic review by two anatomic pathologists with genitourinary cancer expertise for areas of well-defined ccRCC or well-defined sRCC, and areas that had a phenotypical appearance which was intermediate between these well-defined phenotypes (Figure 1A-B).

**Figure 1.**
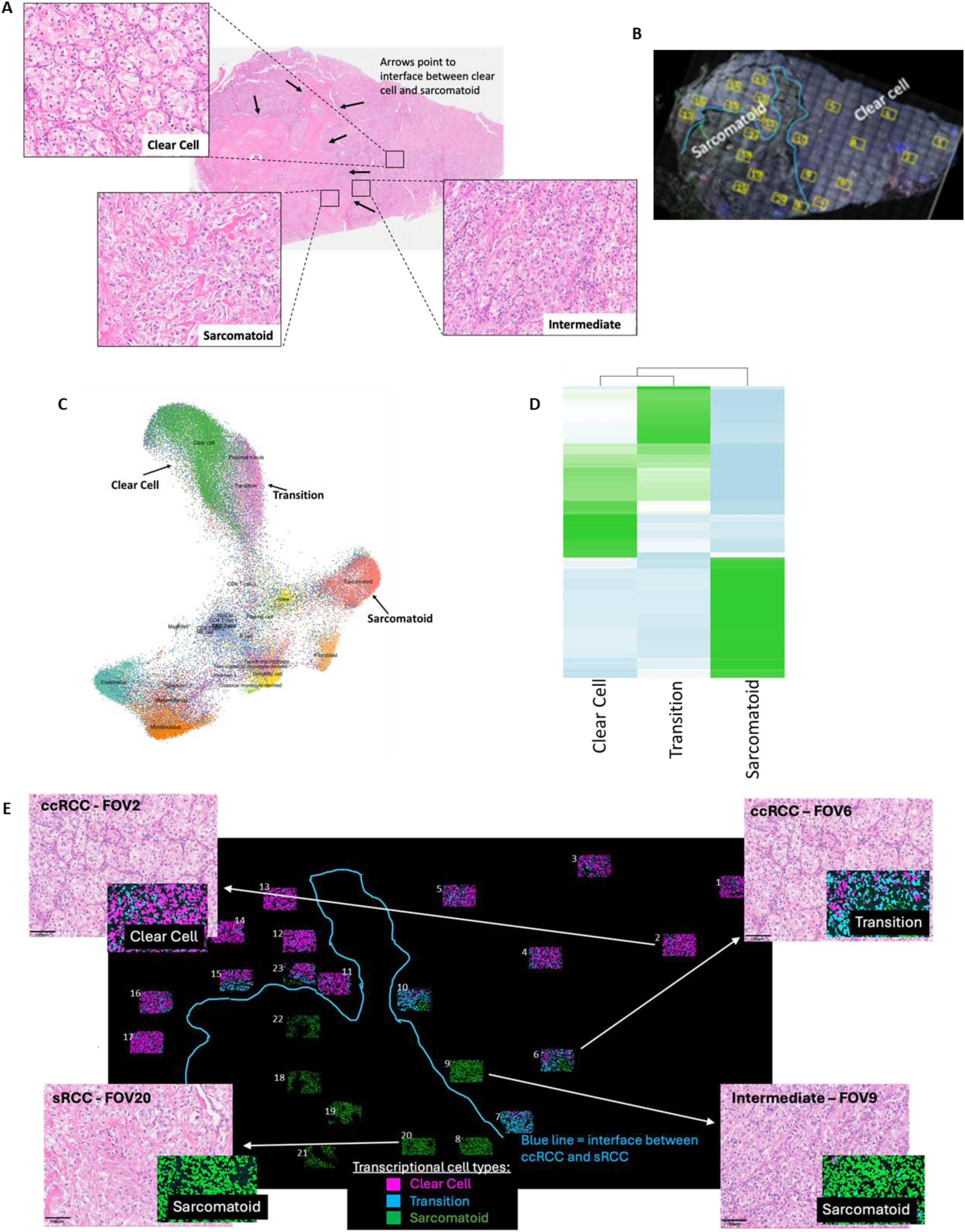
Spatial profiling reveals a unique tumor cell population along the spatial transition between ccRCC and sRCC. (A) Hematoxylin and eosin (H&E) image of sRCC tumor, Sample 1, with arrows pointing to the interface between the ccRCC and sRCC regions and magnified views showing the variation of histologic appearance. (B) Fluorescent image with location of CosMx fields of view (FOVs). The drawn blue line represents the location of spatial interface between clear cell and sarcomatoid for reference. (C) Uniform manifold approximation and projection (UMAP) of malignant and non-malignant cells captured across all FOVs from Specimen 1. (D) Heat map showing comparative gene expression of the tumor cell populations. (E) Spatial mapping of tumor cell types identified by gene expression, colored as indicated in legend. Four representative FOVs are highlighted showing H&E of the FOV with corresponding transcriptional mapped cell types.

We captured approximately 14 million transcripts from 60,000 single cells and performed semi-supervised clustering which was projected in UMAP space (Figure 1C; high definition UMAP in Figure S1). Multiple immune cell populations, several stromal cell populations and three tumor cell populations were identified. Evaluation of the 3 tumor populations demonstrated differences in gene expression (Figure 1D) demonstrating that these are distinct cell populations. We mapped these three annotated cell populations back to the tissue used for selecting FOVs (Figure 1E). This included one tumor cell population, shown in pink (Figure 1E) which mapped to histologically ccRCC areas and was defined as “Clear Cell”, and another population shown in green which mapped to sarcomatoid areas and was defined as “Sarcomatoid” (Figure1E). A third tumor cell population, colored blue (Figure 1E), mapped to histologically clear cell regions approaching the spatial interface of ccRCC to sRCC. Because of its spatial location, we hypothesized the blue cell population may represent a transition state from clear cell to sarcomatoid, and we labeled it “Transition”. Going forward, capitalized “Clear Cell”, “Sarcomatoid” and “Transition” will refer to the transcriptional state and lower case “clear cell” or “ccRCC” or “sarcomatoid” or “sRCC” or “intermediate” will refer to the histologic state.

Histology from hematoxylin and eosin (H&E) of the adjacent cut slide was matched to the spatial transcriptomic data (Figure 1E). Most histologically ccRCC areas had a majority transcriptional cell type of Clear Cell with sparse intermixing of Transition cells. All histologically sarcomatoid FOVs had concordant Sarcomatoid transcriptional makeup. However, FOVs near the spatial interface between ccRCC and sRCC had unique findings. Three FOVs (FOV 6, 7, 10) in histologically-defined clear cell areas nearest the interface had a predominant transcriptomic makeup of Transition (Figure 1E), yet all these FOVs contained tumor cells that were morphologically indistinguishable from other clear cell areas. FOV 9 contained cells that histologically appeared to be intermediate between ccRCC and sRCC, with some clear cell features but elongated cell shape and a loss of the overarching nested structure. Despite morphologic differences in FOV9 compared to distinct sRCC regions, cells in FOV9 expressed a fully Sarcomatoid transcriptional pattern (Figure 1E). These results suggest spatial transcriptomics can identify progressive transcriptomic changes prior to morphologic changes.

### Transition cells exist in a hybrid EMT state along a continuum from clear cell to sarcomatoid

Since sRCC is thought to arise through EMT and EMT occurs on a continuum, we evaluated whether Transition cells exist in a hybrid EMT state between Clear Cell and Sarcomatoid. We created an annotated list to designate epithelial or mesenchymal associated genes within the original 960-gene panel (Table S2) based on a published RCC-specific EMT signature (25) and several other seminal EMT signatures (26, 27, 28). Differential gene expression between Clear Cells and Transition Cells (Figure 2A) shows Clear Cells have higher expression of epithelial-related genes including canonical epithelial marker *CDH1*, or E-cadherin. Transition cells had differentially higher expression of mesenchymal genes including two genes, *COL3A1* and *COL6A2* which encode for collagen proteins, notable because sarcomatoid cells are known to exhibit abnormally high levels of collagen synthesis (29), demonstrating gain of sarcomatoid-like features in the Transition cell state. We performed gene set enrichment analysis (30) based on Hallmark pathways(31) with incorporation of our annotated epithelial and mesenchymal gene list and found the epithelial signature was the most enriched pathway in Clear Cell compared to Transition, and the mesenchymal signature and hallmark EMT pathway were among the most enriched in Transition compared to Clear Cell (Figure 2B). We repeated similar analysis between Transition and Sarcomatoid cells and found Transition cells differentially express more epithelial genes compared to Sarcomatoid, and Sarcomatoid cells express more mesenchymal genes than Transition cells (Figure 2C). GSEA between Transition and Sarcomatoid demonstrates the hallmark EMT pathway as the most upregulated pathway in Sarcomatoid compared to Transition (Figure 2D). This supports the hypothesis that the Transition cell type is a distinct state along a continuum from ccRCC to sRCC where Transition cells express both epithelial and mesenchymal features. Of note, other than the EMT hallmark pathway and mesenchymal gene set, the 3 other top upregulated pathways in Sarcomatoid cells were E2F targets, G2M checkpoint, and UV response, all pathways previously reported to be increased in sRCC (13, 32) (Figure 2D). However, inflammatory related pathways, including inflammatory response, TNFA signaling, and IL6 JAK STAT3 signaling, which previously have been implicated in sRCC (13, 32), were upregulated in the Transition cells compared to Sarcomatoid cells (Figure 2D).

**Figure 2.**
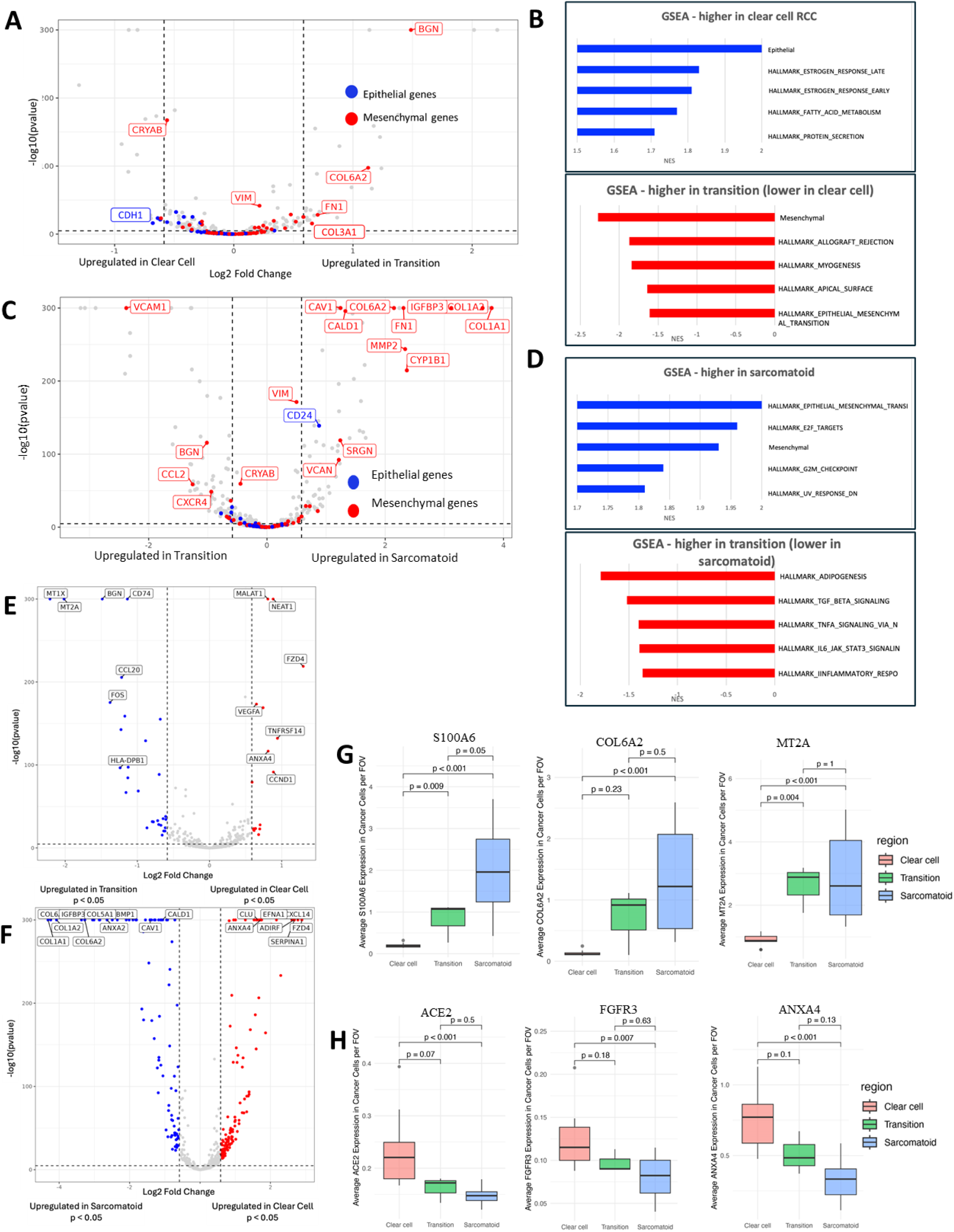
Transition cells exist in a hybrid EMT state along an EMT continuum from clear cell to Sarcomatoid. (A) Differential gene expression (DGE) of Clear Cell versus Transition with highlighted genes from the annotated epithelial (blue) or mesenchymal (red) gene list. (B) Gene set enrichment analysis (GSEA) between Clear Cell and Transition including annotated epithelial and mesenchymal list and Hallmark pathways. (C) DGE of Transition versus Sarcomatoid with highlighted genes from annotated epithelial (blue) or mesenchymal (red) gene list. (D) GSEA between Transition and Sarcomatoid including annotated epithelial and mesenchymal list and Hallmark pathways. (E) DGE (all genes) between Clear Cell (red) and Transition (blue). (F) DGE between Clear Cell (red) and Sarcomatoid (blue). (G) Plots of average gene expression computed from single cell level and averaged per FOV across tumor region of genes mutually increased from Clear Cell to Transition and Sarcomatoid. (H) Plots of average gene expression computed from single cell level and averaged per FOV across tumor region of genes mutually decreased from Clear Cell to Transition and Sarcomatoid.

We hypothesized that if the Transition cells exist along a continuum from Clear Cell to Sarcomatoid, there would be several genes with a continuous increase or decrease along the transformation. To test this, we evaluated differential gene expression between Clear Cell and Transition, now examining all genes rather than just EMT-related genes, (Figure 2E) and between Clear Cell and Sarcomatoid clusters (Figure 2F). We evaluated genes that were mutually increased or decreased in both Transition and Sarcomatoid compared to Clear Cell as the base. We found 7 genes upregulated in both Transition and Sarcomatoid compared to Clear Cell. We evaluated the single cell expression of these genes in tumor cells of regions designated as Clear Cell, Transition, or Sarcomatoid based on majority tumor cell type. Although not all showed significance, likely due to a mixture of cells types diluting the effects, 5 of the seven showed a continuous trend towards increase from Clear Cell to Transition to Sarcomatoid regions, showing there are gradient gene changes in a spatial pattern in this tumor (select genes shown in Figure 2G, expanded list in Table 1, full list Table S3). Two of the 7 genes upregulated in both Transition and Sarcomatoid, *COL6A2, COL3A1*, are related to collagen synthesis as previously mentioned and four are related to EMT (*FN1, S100A6, MT1X, MT2A)*(33, 34, 35, 36, 37). Interestingly, the 2 genes upregulated in both Transition and Sarcomatoid, but with a greater fold change in Transition and trend towards higher expression in Transition regions, *MT1X* and *MT2A*, are associated with hybrid or partial EMT states (36, 37), which may explain their increased expression in the Transition state (Figure 2G, Table 1). We identified 14 genes down regulated in both Transition and Sarcomatoid and plotted the single cell gene expression in tumor regions as above. 12 of the 14 genes showed a significant or trend towards decrease expression from Clear Cell to Transition to Sarcomatoid regions (select genes shown in Figure 2H). Genes downregulated in a gradient manner from Clear Cell to Transition to Sarcomatoid included *FGFR3* involved in cell growth and development (38), and *ANXA4* responsible for modulating cell membrane permeability and cell growth (39) amongst others. Over all, these analyses demonstrated Transition cells exist in a hybrid EMT state between Clear Cell and Sarcomatoid, and that there are several genes with a continuous increase or decrease from Clear Cell to Transition to Sarcomatoid, lending evidence to the Transition state being a stable state along a continuum of transformation from ccRCC to sRCC.

**Table 1.**
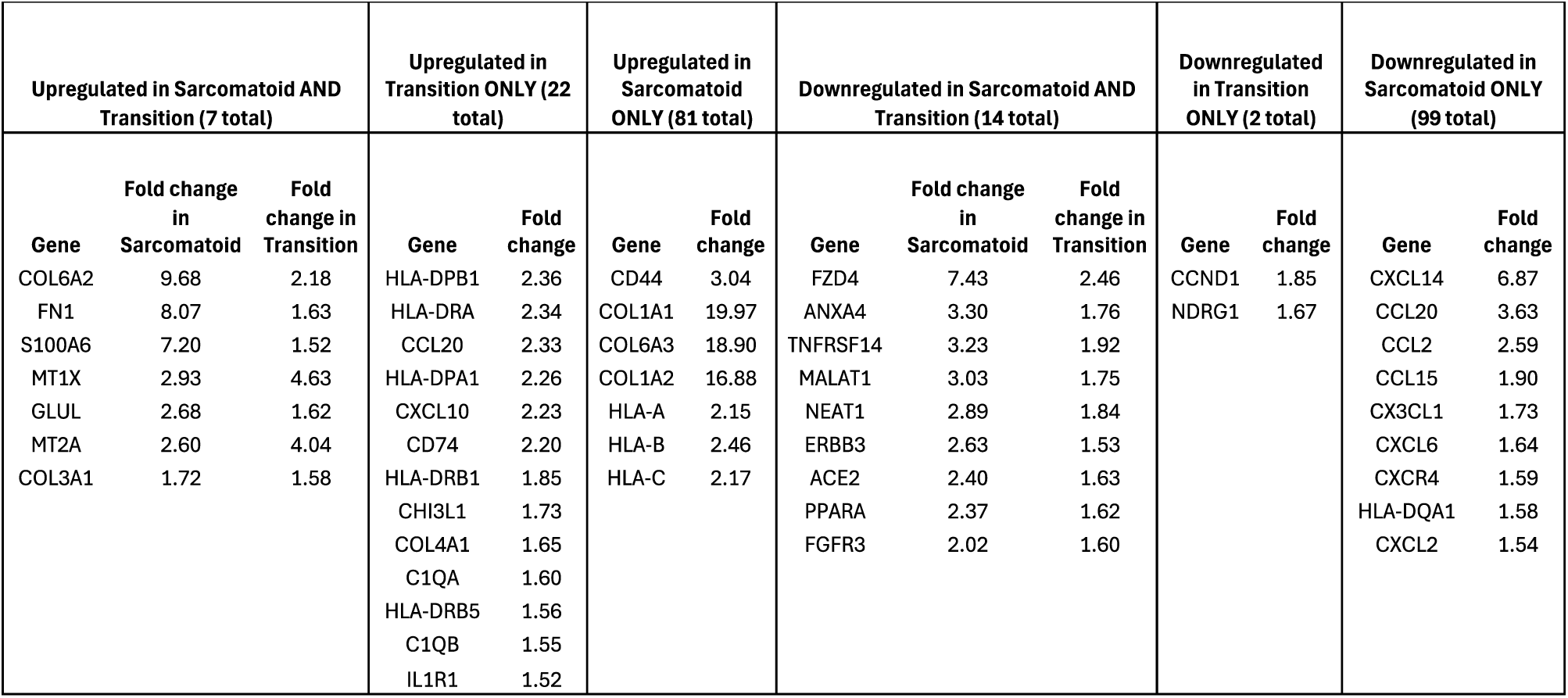
Table showing selected differential gene changes in Transition cells, Sarcomatoid cells, or both in comparison to Clear Cells as the baseline (full gene list available in supplemental Table S3). Transition cells have an increase in cytokines and other factors responsible for inflammation, while sarcomatoid cells have decreased cytokines and inflammatory factors but increased EMT and stemness-related gene expression. All fold changes are listed in the table and all p values are < 0.05.

### Unique tumor immune microenvironments exist as clear cells progress to sarcomatoid, which may be largely shaped by the Transition state

We examined the differential gene expression analysis between Clear Cell and Transition and between Clear Cell and Sarcomatoid (Figure 2E-F) and noted that the majority of the genes significantly upregulated only in Transition were related to inflammation and immune cell recruitment (Table 1). For example, *CCL20*, a well-known chemokine for various immune cells (40), *CXCL10* which mediates immune response and recruitment of T cells (41), *IL1R1*, a potent pro-inflammatory receptor (42), and *CHI3L1*, which activates and recruits macrophages and is also implicated in stimulating macrophage PD-L1 expression (43, 44). We also saw an increase in MHC Class II factors, which are known to shape immune response and tumor response to immunotherapy (45) as well as *CD74*, an invariant chain involved in the formation and transport of MHC II peptides. These findings suggest an increase in inflammatory and immunogenic properties as RCC cells undergo this transition. If Sarcomatoid cells were directing the immune infiltrate in sRCC tumors, we would expect an upregulation of many immune-related genes in Sarcomatoid cells as well. However, of the genes upregulated only in Sarcomatoid cells, the only primarily immune-related genes were MHC Class I factors *HLA-A*, *HLA-B*, and *HLA-C,* which is notable since increased MHC Class I factor expression correlates with immunotherapy response (46). Nearly all the other genes upregulated in Sarcomatoid alone were related to EMT, stemness, and other pathways of tumor progression (Table S3). In fact, many cytokines and chemokines were uniquely downregulated in Sarcomatoid cells (Table 1), suggesting Sarcomatoid cells may actually have decreased ability to recruit immune cells. This supports a hypothesis that the Transition cell state may drive immune infiltrate to sRCC tumors, and that increased MHC Class I factors on Sarcomatoid cells may contribute to immunotherapy response.

To understand whether the immune properties of the Transition state led to changes in the immune cell component of the tumor microenvironment, we evaluated the associated immune microenvironment of Transition regions in comparison to Clear Cell or Sarcomatoid regions based on the majority of transcriptional tumor cell type within each FOV. We observed a shift in immune cell makeup from Clear Cell to Transition and Sarcomatoid (Figure 3A). Interestingly, although Clear Cells and Transition cells were far more similar to each other transcriptionally than either were to Sarcomatoid cells (Figure 1D), the immune microenvironment of Transition and Sarcomatoid were very similar to each other and markedly different from Clear Cell (Figure 3A). Most notably, tissue macrophages were far more prevalent in Transition and Sarcomatoid areas compared to Clear Cell areas (Figure 3A-B). Cell-cell proximity analysis (Figure 3C) showed macrophages are more spatially colocalized with Transition and Sarcomatoid cells compared to Clear Cells. This was confirmed when we mapped macrophages back to the original histological sections (Figure. 3D). To determine if there were macrophage subpopulations, we performed sub-clustering; however, this did not result in distinct subpopulations (not shown), albeit a top marker for these macrophages was CD163, a well-established M2-like marker.

**Figure 3.**
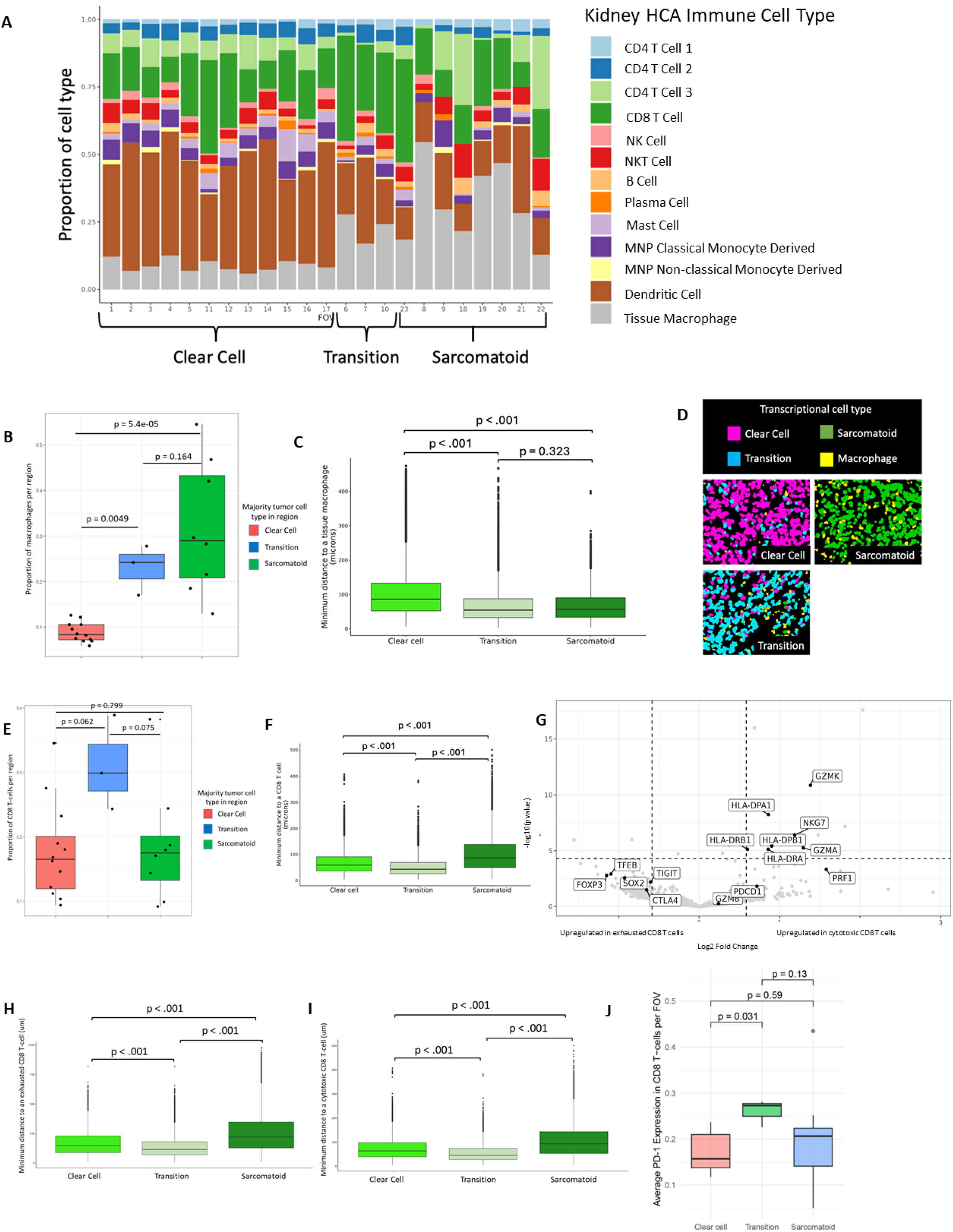
Unique immune microenvironmental changes exist between areas of Clear Cell, Transition and Sarcomatoid. (A) Bar plot showing the respective proportion of each color-coded immune cell type in each FOV (each column represents an FOV). FOVs are ordered by majority tumor cell type makeup (Clear cell, Transition, or Sarcomatoid). (B) Box plot of proportion of macrophages per FOV in FOVs with predominantly Clear Cell, Transition, or Sarcomatoid as the majority tumor cell type. (C) Minimum cell to cell distance from each tumor cell type to nearest macrophage. (D) Representative regions of mapped cell types in a Clear Cell, Transition, and Sarcomatoid region showing macrophage content amongst tumor cells. (E) Box plot of proportion of CD8 T-cells per FOV in FOVs with predominantly Clear Cell, Transition, or Sarcomatoid as the majority tumor cell type. (F) Minimum cell to cell distance of each tumor cell to nearest CD8 T cell. (G) DGE between exhausted and cytotoxic CD8 T-cell clusters. (H) Minimum cell to cell distance from each tumor cell type to nearest exhausted CD8 T-cell (H) or cytotoxic CD8 T-cell (I). (J) Boxplot of average PD-1 expression in CD8 T-cells computed from single cell level and averaged per FOV across each tumor region.

CD8 T-cells were present throughout the tumor but we noticed an increased abundance in Transition FOV’s (Figure 3A, E). CD8 T-cells also showed greater colocalization with Transition cells than other tumor cells in cell-cell proximity analysis (Figure 3F). Sarcomatoid cells were least spatially associated with CD8 T cells (Figure 3F). Supervised sub-clustering of CD8 T-cells split these cells into two main populations. One CD8 T-cell cluster showed differentially higher levels of cytotoxic markers including *GZMA, GZMK,* and (although not reaching significance) *PRF1,* leading us to label this cluster as cytotoxic CD8 T-cells (Figure 3G). The second cluster had much lower expression of cytotoxic markers and differentially higher levels of inhibitory receptors and markers of exhaustion including *TIGIT, CTLA4, FOXP3* (although not reaching significance), and we deemed this to represent a more exhausted CD8 T-cell state (Figure 3G). Cell-cell proximity analysis showed both exhausted and cytotoxic CD8 T-cells were more closely associated with Transition cells than any other tumor cell type (Figure 3H-I). In comparing CD8 T-cell clusters, we noted an unexpected increase in PD-1 expression in the cytotoxic T-cells compared to exhausted (although not reaching significance). Since PD-1+ T-cells have implications in immune checkpoint inhibitor response, we evaluated for PD-1 expression at the single cell level in CD8 T-cells based on tumor region. We found that CD8 T-cells in Transition regions expressed higher levels of PD-1 than in Clear Cell or Sarcomatoid regions (Figure 3J). These findings suggest PD-1 directed immune checkpoint blockade may be most highly effective in Transition areas of sRCC tumors and explains why these tumors may be highly sensitive to PD-1 directed therapy.

### Transition cell state is validated in GeoMx dataset of multiple sRCC tumors

The novel findings of a Transition cell state and associated immune changes led us to examine these features spatially in additional sRCC tumors using NanoString GeoMx digital spatial profiling, which generates bulk region of interest expression data rather than single cell-resolved data. Our dataset included an expanded 47 regions of interest (ROIs) from a near adjacent section of the initial RCC tissue, as well as 49 ROIs from two additional RCCs, for a total of 96 distinct regions analyzed.

To discover changes in gene expression between the different transcriptionally-defined cell types, we turned to the differential expression between Clear Cell and Transition cells (Figure 2E, full gene list in Table S3) and between Clear Cell and Sarcomatoid cells (Figure 2F, Table S3). We created gene signatures using differentially expressed genes with p<0.05 and log2fold change>0.6 (Table S4) and applied these gene signatures to the tumor cell segmented regions of our GeoMx dataset. We began with the near adjacent section of the section used in the CosMx analysis, but with an expanded number of ROIs (Figure 4A). As expected, all ROIs annotated as sarcomatoid based on histology had high transcriptional Sarcomatoid scores. Like the CosMx findings, the GeoMx analysis revealed that ROIs with high Transition scores cluster along the interface between clear cell and sarcomatoid. However, two ROI’s (highlighted in Fig 4A as Low PanCK) were present in the middle of a pure clear cell region where we would not have expected to have a Transition signature. Interestingly, on closer analysis, these two ROIs show a notable decrease in PanCK amongst otherwise stronger PanCK staining in clear cell regions of this tumor. In this tumor, we see loss of PanCK staining as the tumor transitions from clear cell to sarcomatoid regions. This is consistent with EMT and sarcomatoid trans differentiation as PanCK is an epithelial marker. Thus, it is possible these two Transition areas may represent an additional focus of the tumor that is in early progression to sarcomatoid.

**Figure 4.**
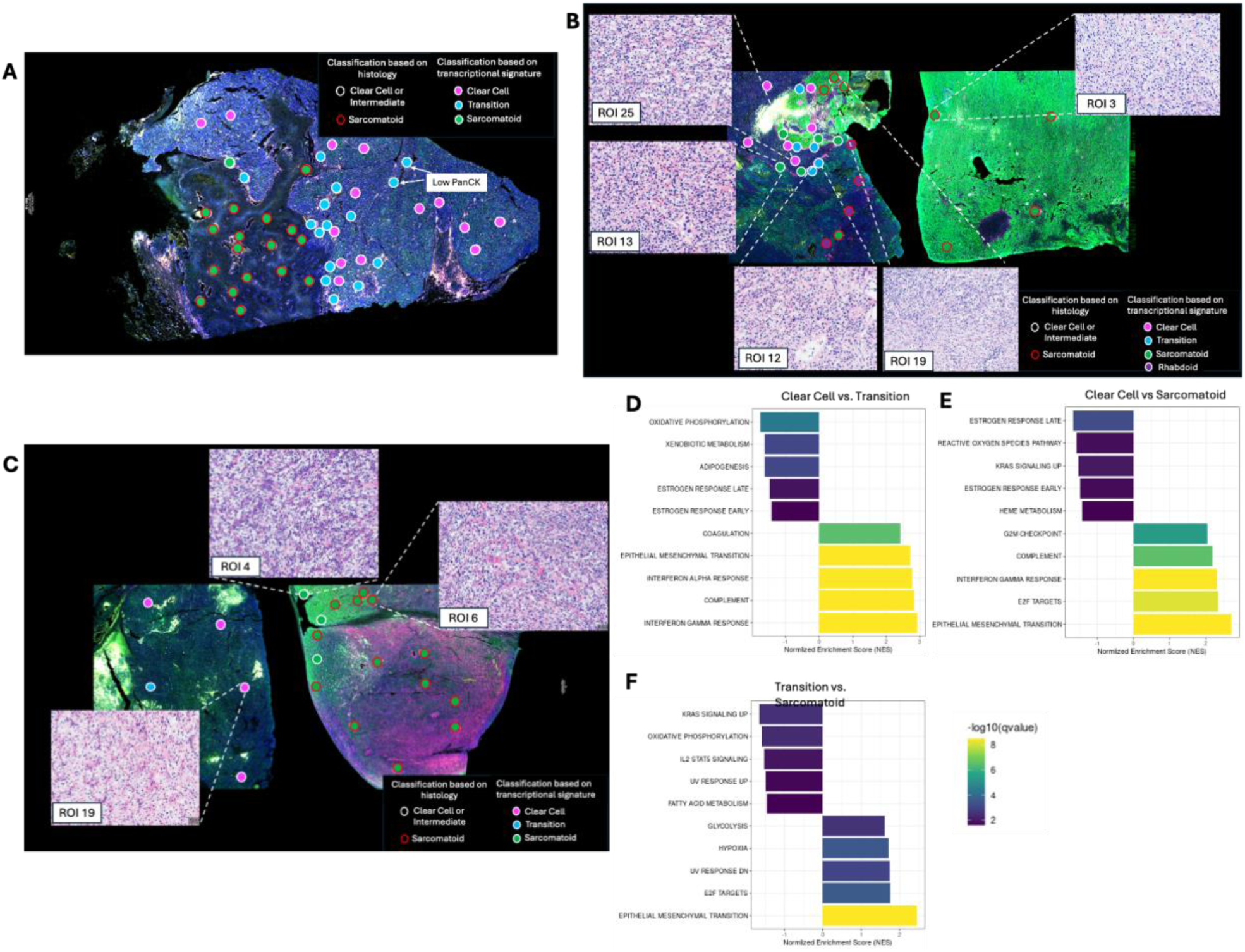
GeoMx analysis in three sarcomatoid tumors shows similar spatial pattern of Transition cell state and transcriptional changes preceding histologic changes. (A) Spatial mapping of GeoMx regions of interest (ROIs) in Specimen 1, where each point represents an ROI, colored by the corresponding transcriptional cell state signature in the legend. The circular outline of each point represents the original histologic annotation where red is sarcomatoid and white is clear cell or intermediate histology. Two regions chosen for low fluorescent expression of PanCK amongst otherwise high expression regions are highlighted. (B) Spatial mapping of histologic and transcriptional ROI annotations in Specimen 2 and (C) Specimen 3 with several representative corresponding H&E areas highlighted. (D) GSEA of Clear Cell vs. Transition regions using Hallmark pathways with +NES = higher in Transition and -NES = higher in Clear Cell and (E) Clear Cell vs. Sarcomatoid with +NES = higher in Sarcomatoid and -NES = higher in Clear Cell and (F) Transition vs. Sarcomatoid with +NES = higher in Sarcomatoid and -NES = higher in Transition.

The same process was applied to two additional RCC tumors. Both tumors had areas of rhabdoid histology as well as sarcomatoid histology. While previous literature has shown that the molecular features of rhabdoid and sarcomatoid do not significantly differ (13), we eliminated rhabdoid areas from analyses to avoid heterogeneity. In one tumor (Figure 4B), we see similar findings to the original tumor, where some histologic clear cell regions express Transition or Sarcomatoid transcriptional pattern. For example, ROI 25 shows clear cell histology and a matching Clear Cell transcription signature but ROI 12, which is along the border between the clear cell and sarcomatoid/rhabdoid region, has a Transition phenotype despite clear cell histology (Figure 4B). Additionally, like the original tumor, some regions that were annotated as clear cell based on histology and are adjacent to sarcomatoid regions have a Sarcomatoid transcriptional pattern, such as ROI 13, despite lacking the full sarcomatoid morphology of sarcomatoid-annotated areas such as ROI 19 and ROI 3 (Figure 4B).

In the final RCC (Figure 4C), we again see a similar pattern where histologically clear cell areas near sarcomatoid areas have a Sarcomatoid transcriptional phenotype, for example, ROI 4.

Comparing the histology of ROI 4 to an ROI with Clear Cell phenotype, ROI 19, we see that cells in ROI 4 appear more elongated and less organized, similar to the “intermediate” histology we previously saw in in the original tumor. Collectively, these findings validate the existence of a spatial transition zone in sRCC and also show that the Transition gene set can be applied to bulk transcriptional data of focal areas, as performed in the GeoMx platform, in RCC.

To evaluate functional changes from Clear Cell to Transition to Sarcomatoid leveraging the full transcriptome, we applied GSEA comparing the ROIs labeled as Clear Cell vs. Transition vs. Sarcomatoid based on their transcriptional gene signature (Figure 4D-F). We validate that even with the full transcriptome, the Epithelial to Mesenchymal pathway is among the most upregulated in Transition compared to Clear Cell (Figure 4D), as well as in Sarcomatoid compared to Clear Cell (Figure 4E) and Sarcomatoid compared to Transition (Figure 4F). We also identify other pathways that were seen in the CosMx analysis, including upregulation of the G2M Checkpoint and E2F Targets in Sarcomatoid regions, validating findings from CosMx and also in agreement with previously identified pathways in sRCC (Figure 4E-F) (13).

### Multiplex immunofluorescence demonstrates macrophages are associated with mesenchymal areas consistent with Transition and Sarcomatoid regions and clustering of CD8 T-cells along mesenchymal borders

To determine if the immune microenvironment changes that were observed in the CosMx analysis were present across a larger cohort of sarcomatoid tumors we performed multiplex immunofluorescent staining on 29 sRCC specimens, using antibodies to CD3 (T-cell), CD8 (CD8 T-cell), CD163 (M2-like macrophage), PD-L1, E-cadherin (epithelial marker) and N-cadherin (mesenchymal marker).

In general, N-cadherin staining was high in sarcomatoid areas and lower in clear cell areas, consistent with the expected EMT pattern. N-cadherin staining was present at intermediate levels in clear cell areas spatially near sarcomatoid areas of several tumors (Figure 5A), supporting our prior findings of a hybrid EMT transition state spatially near sRCC. E-cadherin staining did not follow a predictable pattern, consistent with some literature suggesting E-cadherin does not always change with EMT (47, 48), and therefore, we elected to use high N-cadherin as the main designation of mesenchymal/sarcomatoid areas and low N-cadherin as epithelial/clear cell areas. We found a strong correlation between N-cadherin and CD163 (Figure 5B-C), corroborating our prior findings of M2-like macrophage association with sRCC regions.

**Figure 5.**
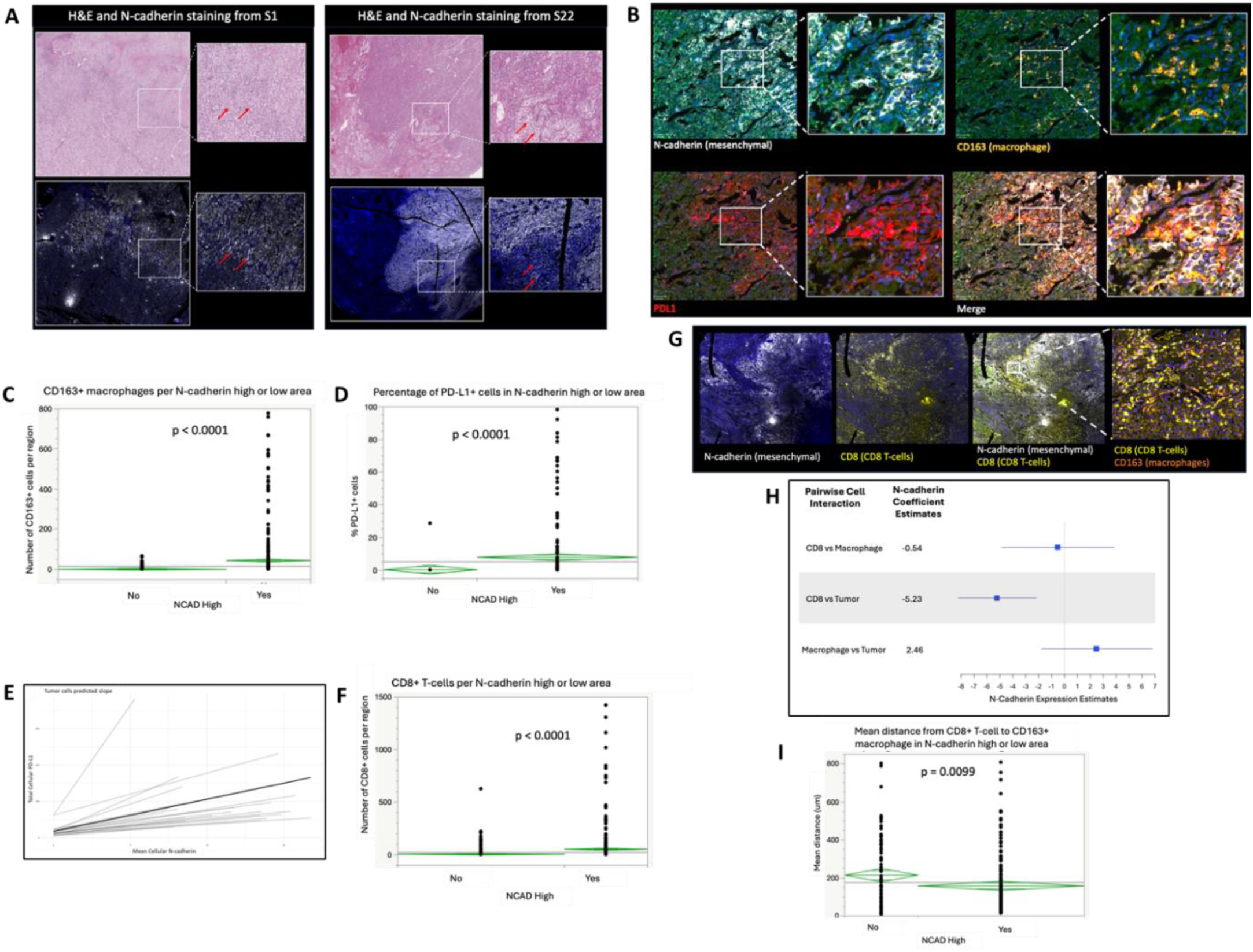
Immune microenvironment changes that occur during progression from clear cell to sarcomatoid are validated in multiplex immunofluorescent (mIF) staining of 29 specimens. (A) Representative images from mIF of 29 specimens from 20 sRCC tumors showing spatial distribution of mesenchymal marker, N-cadherin, staining (white) with corresponding H&E. (B) Representative images showing colocalization of N-cadherin (white), PD-L1 (red), and CD163 (orange) staining in sRCC areas. (C) Plot of number of CD163+ cells in N-cadherin high versus N-cadherin low areas. (D) Plot of percentage of PD-L1 positive cells in N-cadherin high versus low areas. (E) Patient specific (grey) and average (black) regression lines for association between mean N-cadherin and total cellular PD-L1 expression in tumor cells. (F) Plot of number of CD8 T cells in N-cadherin high versus low areas. (G) Representative area showing pattern of CD8 staining seen in many tumors where CD8 T cells localize to outer edge of N-cadherin high areas. (H) Plot of the effect of N-cadherin expression on tumor cell, macrophage, and CD8 T cell colocalization. A positive value indicates two cells are closer together as N-cadherin increases and negative indicates the two cells are more separated as N-cadherin increases. (I) Mean distance between CD8+ cells and CD163+ cells in N-cadherin high versus low areas compared to

PD-L1 has previously been shown to be increased in the sarcomatoid components of sRCC tumors (49), and we sought to evaluate the spatial relationship of PD-L1 expression in relation to N-cadherin expression, since EMT has been reported to upregulate PD-L1 expression (50). We found strong association between PD-L1 expression and N-cadherin expression (Figure 5B, D). We developed a linear mixed model, accounting for between patient heterogeneity, and this showed a positive association between N-cadherin and PD-L1 expression in tumor cells (p<0.05), suggesting PD-L1 expression may be directly related to N-cadherin expression (Figure 5E). Additionally, PD-L1 expression was identified not only on tumor cells, but also on macrophages in N-cadherin high areas (Figure 5B).

Evaluation of CD8 T cell spatial patterns revealed that CD8 staining was higher in N-cadherin high regions across all 29 sRCC specimens (Figure 5F), however closer visualization of the staining patterns showed several tumors in which CD8 T cells aggregated along the outer rim of N-cadherin areas (Figure 5G). To better quantify this, we statistically modeled the association between spatial colocalization of CD8 T cells, macrophages, and tumor cells with corresponding changes in N-cadherin using spatial analytical tools across all 29 sRCC specimens. We found as N-cadherin expression increased, CD8 T cells and tumor cells tend to spatially separate, and this association is statistically significant (Figure 5H). On the other hand, macrophages and tumor cells colocalize more as N-cadherin increased (although not reaching statistical significance, Figure 5H). This shows the importance of the single cell spatial resolution because despite increased CD8 T cell density in sRCC areas, the T cells and tumor cells become less colocalized, suggesting decreased interaction between them.

In the CosMx data we noted several instances of PD-L1+ macrophages and PD-1+ CD8 T cells appearing to colocalize (Figure S2), which led us to evaluate whether there was a spatial association between macrophages and CD8 T-cells in the full cohort of sRCC tumors. We found CD8 T-cells and macrophages were more spatially associated in N-cadherin high areas than in N-cadherin low areas (Figure 5I) on a binary measurement, although this finding did not remain statistically significant when evaluating over the continuum of N-cadherin expression (Figure 5H).

Overall, multiplex immunofluorescent staining in this cohort of 29 sRCC specimens validated the CosMx findings and, among other findings, found that M2-like macrophages are strongly associated with sarcomatoid and EMT-high clear cell areas.

### External data set validation shows the Transition state is found in clear cell regions of sarcomatoid tumors, metastatic tumors, and in pure ccRCC tumors with poor outcomes

To validate the existence of the Transition cell state and its correlation with M2-like macrophages, we turned to an external single cell spatial transcriptomic dataset from Moffitt Cancer Center (51). This dataset included the same 960 plex NanoString CosMx analysis of a tissue microarray (TMA) composed of 21 pure ccRCC or ccRCC/sRCC specimens (primaries or metastases). Transition scores were calculated for each tumor cell following the same methodology as the prior GeoMx analysis, but now at the single cell level. The median Transition score was significantly higher in clear cell regions of tumors that had sRCC diagnosis (Clear cell; Primary: Sarcomatoid) than of pure ccRCC tumors (Clear cell; Primary: non-Sarcomatoid) (0.79 vs. 0.73, p=.04; Figure 6A), validating that these transcriptional changes are found in histologically defined clear cell regions of sarcomatoid tumors. Taking advantage of the single cell and spatial resolution of this dataset, we evaluated the association of M2-like macrophages with regions of high transition score. As seen in our prior data, CD163+ macrophages were strongly spatially associated with Transition cells in the clear cell regions of pure ccRCC and sRCC tumors (Figure 6B-C).

**Figure 6.**
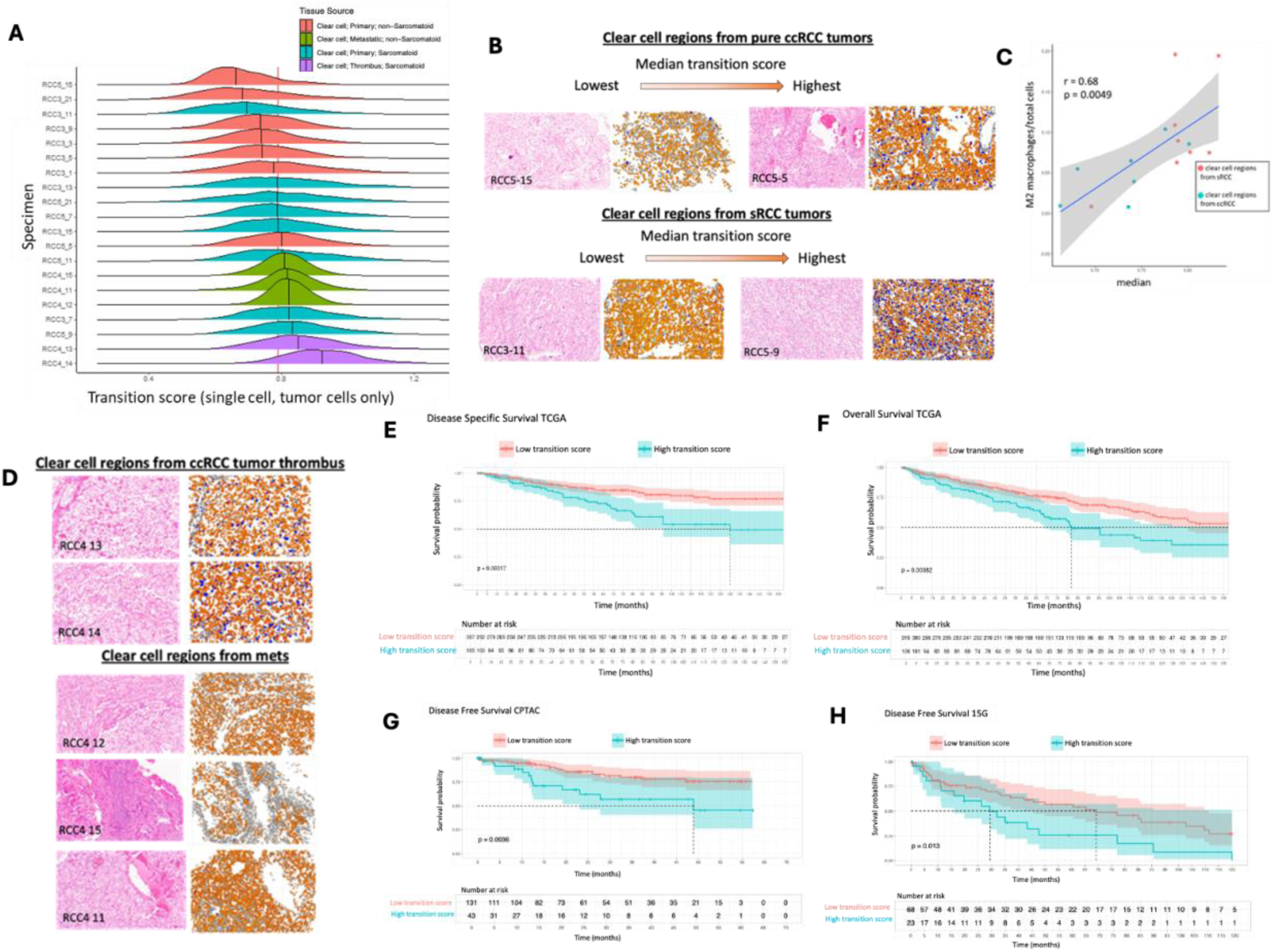
Transition findings are validated in external dataset of 20 tumor with single cell spatial transcriptomics showing spatial correlation with M2-like macrophages, and in large datasets showing correlation with clinical outcome. (A) The “Transition” gene score was applied to a cohort of 20 single cell spatial transcriptomic specimens in tissue microarray (TMA) form. Each curve represents the distribution of transition signature score of all single tumor cells within each TMA core, color coded as indicated in legend. (B) Images of the matched histology and single cell plotting of tumor cell “Transition” score (heat gradient) and CD163+ macrophage infiltrate in blue. The range of lowest to highest median “Transition” score in pure primary ccRCC tumors (top) and ccRCC regions from tumors with sRCC (bottom) is shown with corresponding CD163+ macrophage infiltrate. (C) Correlation between Transition score and M2 macrophage content in ccRCC and sRCC tumors. (D) Images of matched histology, tumor cell Transition score, and M2 macrophages in metastatic and tumor thrombus lesions. (E-H) Kaplan Meier curves showing clinical outcomes for tumors with high Transition score (top quartile) versus low (remaining bottom 75%) for disease specific survival (DSS) in the TCGA (E) and (F) overall survival (OS) in the TCGA, and (G) Disease free survival (DFS) in the CPTAC dataset and the 15G cohort (H).

The Moffitt dataset also included 3 specimens from metastatic lesions and 2 from tumor thrombi, which all had very high Transition scores (Figure 6A, D), supporting the notion that EMT biology in ccRCC tumors leads to aggressive phenotypes. Notably, the 3 metastatic lesions were from patients with pure ccRCC, without known sRCC diagnosis, so the finding of high Transition gene expression in these tumors suggested a transition may be detectable in aggressive tumors which have not yet demonstrated sarcomatoid histology. To explore this further, we utilized several large publicly available datasets of ccRCC tumors without any known sRCC diagnosis, including the KIRC TCGA dataset (52), the CPTAC dataset (53), and the 15G dataset (54). We found increased expression of the Transition gene set was associated with worse overall survival (OS) and disease specific survival (DSS) in the TCGA data, and poorer disease-free survival (DFS) in the CPTAC and 15G cohorts (Figure 6 E-H).

This confirms the biologic relevance of the Transition transcriptional changes, indicating that these changes contribute to tumor aggressiveness and poor prognosis, and that such changes may be identifiable in ccRCC tumors prior to development of full sarcomatoid changes.

### M2-like macrophages induce EMT, PD-L1, and other sarcomatoid-like changes in ccRCC cells through SPP1

Since there was a strong association of M2-like macrophages with Transition and Sarcomatoid areas in our spatial data, we explored for functional interactions between ccRCC cells and M2-like macrophages. The CosMx analyses revealed *CCL20* to be highly upregulated in Transition cells compared to Clear Cells. Taken together with the previous reports that tumor cell-derived *CCL20* plays a role in recruitment of myeloid cells and polarization of macrophages towards M2-like phenotype (55, 56) we explored for functional relations between RCC cells lines, CCL20 and macrophage-like cell lines. We hypothesized that CCL20 is upregulated during the EMT transition and contributes towards shaping the unique immune microenvironment during sarcomatoid transition. To test this, we induced EMT in the 786-O ccRCC cell line through exposure to TGFβ, a known EMT inducer (EMT shown to be induced in Figure S3), and found that CCL20 expression was upregulated during EMT (Figure 7A) and the same was found in a second RCC cell line, UOK-127 (Figure S4).

**Figure 7.**
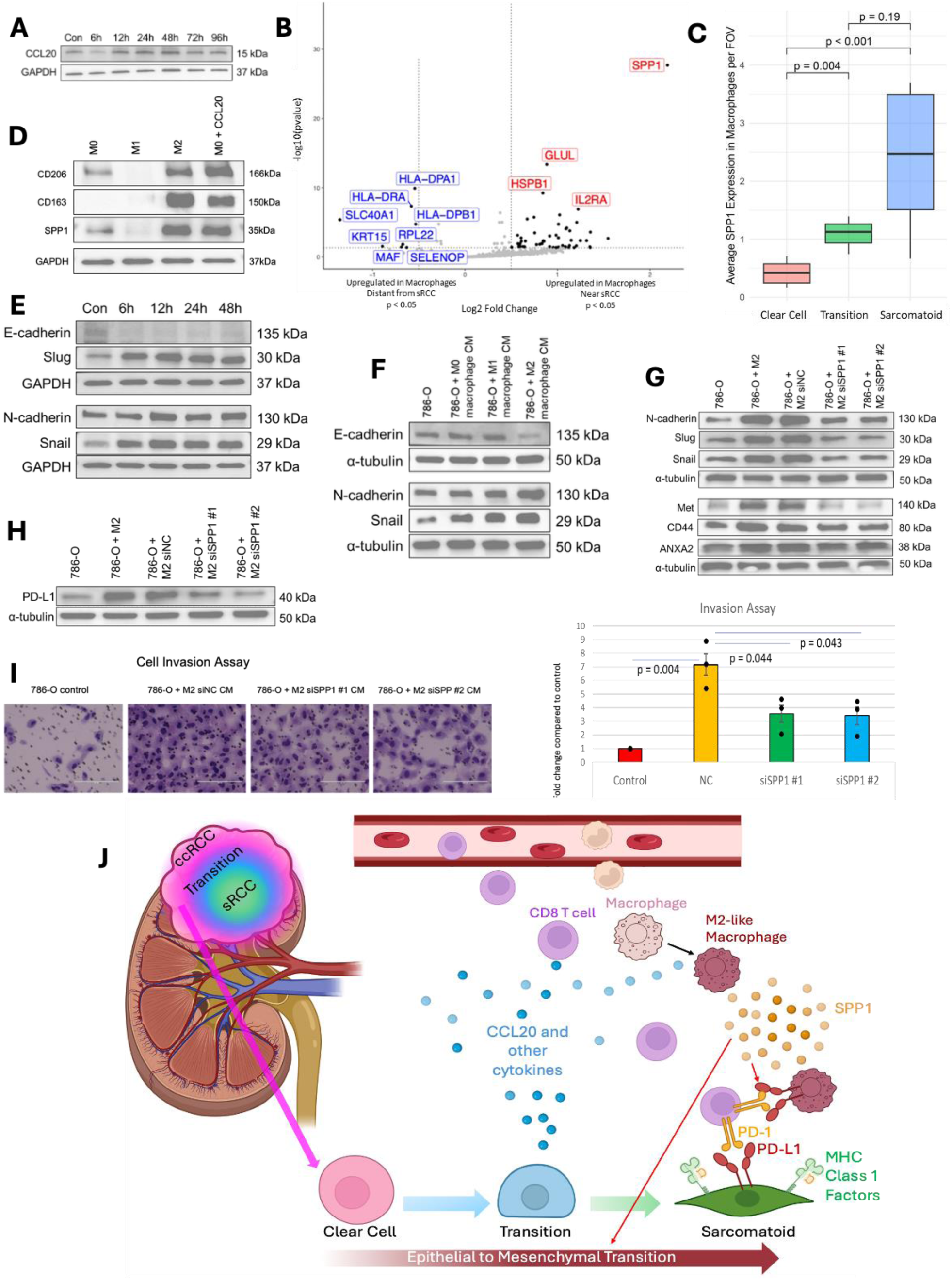
Transition tumor cells polarize macrophages to M2 state via CCL20 and M2 macrophages induce sarcomatoid changes in RCC cells via SPP1. (A) CCL20 protein levels in 786-O RCC cells after induction of EMT by TGFb. (B) DGE between macrophages within 50um of sRCC cells (right) vs >50um (left) from CosMx analysis. (C) Box plot of average SPP1 expression of macrophages in Clear Cell, Transition, or Sarcomatoid FOVs from CosMx analysis. (D) Macrophage differentiation marker and SPP1 protein expression in polarized THP-1 derived macrophages and M0 macrophages treated with recombinant CCL20. (E) EMT protein markers from 786-O cells treated with recombinant SPP1. (F) EMT markers in 786-O cells treated with conditioned media from M0, M1, or M2-like macrophages. (G) EMT marker proteins and several proteins found to be highly differentially expressed in Sarcomatoid cells at the mRNA level (CD44, Met, ANXA2) in 786-O cells treated with conditioned media from M2 macrophages with or without SPP1 knockdown by siRNA. (H) PD-L1 protein expression in 786-O cells treated with conditioned media from M2 macrophages with or without SPP1 knockdown by siRNA. (I) Cell invasion assay representative images and quantification in 786-O cells treated with conditioned media from M2 macrophages with or without SPP1 knockdown by siRNA. (J) Schematic of proposed mechanistic pathway of sarcomatoid transformation. Clear cells undergo transformation to “transition” phenotype in which they express increased cytokines including CCL20 which attract immune infiltrate including macrophages. CCL20 and other cytokines lead to M2-like polarization of macrophages with increased SPP1 expression. SPP1+ macrophages then feedback to tumor cells, inducing EMT and PD-L1 expression and leading to full sarcomatoid-like changes.

Macrophages have been shown to induce EMT in cancer cells, including RCC cells (57), however the mechanism behind this is not well described. To identify potential mechanisms, we analyzed differential gene expression in macrophages near (within 50um) versus distant to Sarcomatoid cells in the CosMx data. This revealed several differentially expressed genes (Figure 7B), but one in particular, *SPP1*, was far more highly expressed in macrophages near sarcomatoid cells compared to those distant to sarcomatoid cells with a greater than 4-fold change (Figure 7B). SPP1, also known as secreted phosphoprotein 1 or osteopontin, has recently been implicated in EMT processes in other disease states (58, 59). We evaluated the gradient of *SPP1* expression at the single cell level in macrophages of predominantly Clear Cell versus Transition versus Sarcomatoid areas and found a strong increase in *SPP1* expression as the tumor cells undergo sarcomatoid transformation (Figure 7C). To determine whether the Transition state may crosstalk with macrophages to increase SPP1 expression through CCL20, we utilized a THP-1 derived macrophage model to evaluate SPP1 expression in M0, M1, and M2 polarized macrophages as well as M0 macrophages treated with recombinant CCL20. We found that recombinant CCL20 increases M2 polarization of macrophages as well as SPP1 expression in macrophages (Figure 7D). Additionally, we observed that classically M2 polarized macrophages also express SPP1 (Figure 7D).

Since *SPP1* encodes for a secreted protein, we explored the effect of recombinant SPP1 on RCC cells in vitro. SPP1 diminished E-cadherin expression and induced expression of Slug, N-cadherin and Snail, which is consistent with induction of EMT (Figure 7E) which was also confirmed in a second RCC cell line, UOK127 (Figure S5). We then co-cultured 786-O cells with THP-1 derived M2-like macrophage conditioned media and found that the conditioned media downregulated E-cadherin an upregulated N-cadherin and Snail demonstrating that it promoted EMT in the RCC cells (Figure 7F) and this was also observed in UOK127 cells (Figure S6). We next evaluated the ability of M2-like macrophages to induce other changes in protein expression that were associated with the sarcomatoid phenotype based on the CosMx and proteomic analyses including upregulation of PD-L1, CD44, MET and ANXA2. These proteins were of interest due to their published association with aggressive or sarcomatoid-like variants of RCC (60, 61, 62, 63, 64). M2-like macrophage conditioned media upregulated expression of these sarcomatoid-associated proteins in 786-O cells (Figure 7G-H). To determine if macrophages mediated this effect through SPP1, we knocked down SPP1 in the macrophages (Figure S7). Knockdown of *SPP1* in the macrophages, decreased the ability of M2-like macrophage conditioned media to induce all the sarcomatoid-associated proteins (Figure 7G-H). These results were also observed in UOK127 cells as well (Figure S8).

Finally, since sarcomatoid cells are known to exhibit high capacity for invasion, we tested the ability of M2-like macrophages to induce invasive ability. M2-like macrophage conditioned media induced RCC cell invasion by 7-fold (Figure 7I) and knockdown of *SPP1* in the macrophages diminished this induction by 50%, replicated in UOK-127 cells (Figure S9). These data suggest that macrophages promote the development of sarcomatoid features in RCC through SPP1.

## DISCUSSION

This study presents a unique look into the dynamics of the sarcomatoid transformation in RCC tumors using multiomic single cell spatial approach. By using matched histology and spatial location, we captured cells along a gradient of transformation from clear cell to sarcomatoid RCC and identified a stable, hybrid EMT Transition state. We showed that cells in this Transition state, rather than sarcomatoid cells themselves, express inflammatory factors leading to the high immune infiltrate seen in sRCC tumors. We demonstrated a strong association of M2-like macrophages in areas of Transition and Sarcomatoid and showed mechanistically that M2-like macrophages induce EMT and sarcomatoid-like changes in RCC cells through SPP1 signaling. We propose that Transition tumor cells recruit macrophages to the tumor through CCL20 and other factors, and macrophages then induce further EMT leading to a full sarcomatoid transformation (Figure 7J). Finally, we demonstrated that genes associated with the transition are found in a subset of pure ccRCC tumors and associated with poor prognosis and may identify tumors in early stages of sarcomatoid transformation.

Multiple prior studies have explored the molecular landscape of sRCC, but almost all have done this by comparing sRCC tumors to pure ccRCC tumors or by comparing the epithelioid component of sRCC tumors to the sarcomatoid component in a binary manner (10, 11, 12, 13, 65, 66). Studying the dynamics of sarcomatoid transformation is exceptionally difficult for several reasons. First, the triggers of sarcomatoid transformation are unknown, making it difficult to induce this process in vitro or in vivo. Additionally, there are several limitations to pre-clinical models of RCC, both sarcomatoid and otherwise, which limit such studies (67). This means most of what is known about sRCC comes from the study of patient specimens. Such analyses have been done using bulk tissues that contain a mixture of both clear cell and sarcomatoid areas resulting in the inability to evaluate epithelioid versus sarcomatoid areas and limiting the ability to understand tumor/immune interactions. Some studies have separated clear cell and sarcomatoid regions of individual tumors using micro-dissection (11, 65), but we are unaware of any studies that have done this at a single cell level, which is necessary to distinguish individual tumor and immune cell states. New spatial technologies allowing single cell resolution present an exciting opportunity to study these features.

We approached this study with the hypothesis that if sRCC arises from EMT, we may be able to detect hybrid EMT states, or a gradient of EMT in tumors. This is a well-described studied concept in EMT literature, with multiple studies validating the existence of stable hybrid EMT states (68, 69). Here, we identified a distinct tumor cell population existing in a hybrid EMT state between the epithelioid clear cells and mesenchymal sarcomatoid cells, and existing spatially between the two. Several findings support this cell state being a transition state, including gradient step changes in several genes from Clear Cell to Transition to Sarcomatoid, validation of transcriptional findings and spatial patterns in other sRCC tumors, and consistent findings of these transcriptomic changes preceding histologic changes. While we are unaware of any prior studies demonstrating a transition state between the extremes of clear cell and sarcomatoid, the notion is not unsupported. A prior study by Bostrom et al. stained ccRCC/sRCC tumors for various EMT markers and found that some changes occurred not only in sarcomatoid regions but also in nearby clear cell regions (70). This is consistent with our findings of increased N-cadherin in clear cell areas as the approach the sarcomatoid areas and supports the notion of a spatial continuum. In fact, for many decades, pathologists have described the appearance of blending morphology between clear cell and sarcomatoid cells, sometimes previously termed “early spindle cell changes” (29, 71, 72). Several tumors in our dataset demonstrated histology with areas of what appeared to have intermediate histology, or early spindle cell changes. However, interestingly, these areas rarely demonstrated the Transition transcriptomic profile but were rather found to have a Sarcomatoid transcriptomic profile. The Transition profile was instead isolated to areas of clear cell histology. We believe this demonstrates that transcriptomic changes occur first, driving later morphologic changes.

Importantly, this would suggest that early transcriptomic changes could be detected before a tumor has progressed fully to sarcomatoid, possibly providing an early opportunity to impact patient management. Indeed, we demonstrated Transition state findings in 3 large cohorts of pure ccRCC which were associated with poor prognosis. It is important to note that our Transition gene set was originally derived from the biology of one tumor, and while we find many components of this signature hold true in validation sets, further work is required before proposing this as a prognostic biomarker. However, we believe our findings demonstrate the significance of transitional findings in ccRCC and serve as a crucial starting point to understand novel mechanisms of kidney cancer progression.

Prior studies have identified gene mutations such as in *TP53, CDKN2A, NF2* and others, which may contribute to sarcomatoid development in RCC. However, when epithelioid components are compared to sarcomatoid components within the same tumor, DNA mutations in the two areas are typically very similar (10, 11, 12, 13), suggesting non-mutational factors influence the sarcomatoid transformation. A recent study by El Zarif et al. demonstrated that consistent alterations in gene regulatory element activity are present in sarcomatoid compared to clear cell tumors, identifying key transcription factors such as *FOSL1,* and suggesting epigenetic changes may drive sRCC changes (65). Our findings support the notion that microenvironmental factors, particularly SPP1+ macrophages in the immune microenvironment, induce transcriptional changes that promote the transformation to sRCC.

Multiple studies have shown that sRCC tumors are highly immune infiltrated but also demonstrate high expression of immune checkpoints (13, 49). While some studies suggest sRCC tumor signaling is high in inflammatory pathways such as Interferon Gamma response and tumor necrosis factor alpha (TNFA) signaling (13, 65), it is not clear whether those signals come from the clear cells versus the sarcomatoid cells, or immune cells. By using single cell spatial technologies, we found that immunogenic signaling is driven by cells in transition to sarcomatoid, and the transitional areas harbor most cytotoxic T cells. The multiplex immunofluorescence staining showed that while CD8 T cells are most prevalent in mesenchymal areas, the staining pattern frequently shows CD8 T cells in a dense border along the outer edges of mesenchymal areas, and CD8 T cells become increasingly spatially separated from tumor cells with increasing mesenchymal tumor state. This highlights the importance of careful and complex analyses at the single cell level and demonstrates that bulk or binary analyses may not be fully revealing, perhaps explaining the discrepancies between our data and prior studies. We found Sarcomatoid cells have elevated expression of MHC Class I molecules which may make the Sarcomatoid cells more susceptible to T cell attack. It follows that immune checkpoint inhibitors of the PD-1/PDL-1 axis could be highly effective in Transition and Sarcomatoid cells compared to Clear cells, as the Transition and Sarcomatoid cells have high expression of PD-L1, and a heavily immune infiltrated environment compared to Clear Cells. Thus, once the PD-1/PD-L1 blockade is lifted, the heightened immune response is observed in the Sarcomatoid cells.

Our findings show a strong M2-like macrophage presence in mesenchymal areas. This agrees with multiple recent studies in non-sarcomatoid RCC demonstrating the role of macrophages in driving aggressive tumor features and promoting immunosuppression (73, 74, 75). The role of SPP1+ macrophages in driving cancer progression has been increasingly studied as novel technologies such as single cell spatial transcriptomics have become increasingly available (58, 59). Here, based on in vitro functional studies, we identify SPP1 as an important driver of macrophage-induced changes, including induction of EMT that promotes the transition of ccRCC to sRCC. These results indicate the need for further investigation of the role of macrophages in RCC tumor progression and invites the potential for macrophage targeted therapies in the future.

There are several limitations to this study. Our initial identification of a transition state came from the multi-region study of a single tumor, and while we validated the transcriptional transition state in multiple cohorts, including 3 whole slide sRCC specimens using GeoMx, and 20 additional single cell spatial transcriptomic specimens in the form of a TMA, further work is required to discern which precise features of the Transition state are true across all tumors.

Another limitation was only a 960-plex gene panel was available at the time of our study. Currently, newer technology has expanded these panels to 6000 genes through full transcriptome. This rapidly developing field of technology presents additional opportunities to glean more information from these spatial studies. Another limitation is our methodology of matching transcriptomic or proteomic findings to an adjacent cut stained for H&E. It is possible that an adjacent cut may differ slightly in cellular content, and we required manual visual matching of regions based on location in the tumor and landmarks, which may lead to slight inconsistencies in matching. Additional limitations include the use of several different cohorts which inherently leads to batch effect. The *in vitro* studies are limited by the use of a monocyte derived cell line differentiated to macrophages, which, while commonly used for in vitro study of macrophages, is likely to differ in many ways from true tumor associated macrophages.

In summary, we present a novel use of multi-omic spatial biology techniques to uncover the features of clear cell to sarcomatoid RCC transformation. We identify for the first time a hybrid EMT cell state, spatially abutting sarcomatoid regions, that we believe represents a state of transition from clear cell to sarcomatoid. This provides a unique prospective on the current understanding of sarcomatoid dedifferentiation in RCC and provides a vast avenue for future study. We provide a spatially comprehensive analysis of immune microenvironment changes along the progression from clear cell to sarcomatoid, identifying inflammatory features of the transition state which may recruit immune cells to sRCC tumors. We demonstrate mechanistically that SPP1+ M2-like macrophages contribute to the development of progressive EMT, immune checkpoint expression, and sarcomatoid features in RCC cells. Further work stemming from these novel findings will lead to a better understanding of tumor cell/immune interactions in RCC progression and immunotherapy response, as well as the potential to identify new therapeutic targets in these aggressive tumors.

## METHODS

### Patient Cohort

20 patients with surgically resected sarcomatoid RCC with no prior systemic therapy and available FFPE tissue blocks were identified through retrospective review of nephrectomy specimens at University of Michigan from 2010 – 2021. All human specimens were deidentified to ensure protection of patient data. All use of human tissue and data wase approved by the University of Michigan Institutional Review Board. H&E from FFPE blocks were reviewed by genitourinary pathologists (authors AU and RM) and blocks containing either epithelioid histology, sarcomatoid, or mixed were selected, for a total of 29 blocks. Of these patients, 18 had clear cell RCC as the primary epithelioid component, 1 had papillary RCC, and 1 had chromophobe RCC. Specimen 1 was selected for CosMx single cell spatial transcriptomics. In addition to Specimen 1, additional specimens 2 and 3 were selected for GeoMx analysis, and all 29 specimens were used for multiplex immunofluorescence.

### Validation Cohorts

Single cell spatial transcriptomic data from the Moffitt cohort included a tissue microarray of 42 core FFPE samples from 21 patients(76). Cores containing tumor with prior immunotherapy treatment were excluded as prior treatment may alter transcriptomic makeup. Cores with full sarcomatoid histology were excluded in order to evaluate the clear cell regions only.

The Cancer Genome Atlas (TCGA) cohort is comprised of patients diagnosed with localized ccRCC (pT1-T4 and M0) in the TCGA who had molecular profiling of their primary tumor and associated oncologic outcomes data available and with samples with a library size <1 million reads removed (total n=382). The Clinical Proteomics Tumor Analysis Consortium ccRCC (CPTAC) data was filtered to remove samples without clinical outcomes correlated or with metastasis (n=191). The 15 Gene cohort (15G) included primary untreated ccRCC tumor specimens (n=91).

### CosMx Spatial Molecular Imager gene expression profiling and analyses

Full details of CosMx chemistry and workflow are found in He et al. 2022 (24). Briefly, Specimen 1 (full block FFPE) was sectioned onto a slide and 24 regions were manually selected based on matching with prior selected regions from the adjacent H&E stained slide, including regions of clear cell, regions of sarcomatoid, and regions with intermediate appearing histology. The slide underwent in situ hybridization of 960 mRNA probes using a pre-commercial version of the CosMx Universal Cell Characterization RNA panel.

### Raw image processing and cell segmentation

Raw image processing and feature extraction were performed using the NanoString SMI data processing pipeline and the XYZ location information of each transcript was extracted as previously described (24). Cell segmentation was performed via a previously described pipeline (77, 78).

### Cell clustering and subtyping

Semi-supervised clustering of cells was performed using the Insitutype algorithm (79). Cell typing was performed based on canonical marker gene expression with reference to the kidney human cell atlas (80) and manual refinement to identify tumor cell populations based on gene expression and matched histology. During cell typing, four distinct tumor populations were identified based on their transcriptional profiles. Following histological review of the regions where these populations were localized, we reclassified them as Clear Cell, Transition, and a single Sarcomatoid group formed by merging two of the original populations.

Supervised clustering of T-cells was performed using the immune cell reference proile in the InSituType package (79).

### Differential Expression

Differential expression was performed within and between cell types using a negative-binomial fixed effects model as implemented in the smiDE package (81). For each comparison, the model included expression in surrounding other cell types as a fixed effect and the total counts of the cell as an offset.

### Gene set enrichment analyses (GSEA)

Annotated “epithelial” and “mesenchymal” gene lists were manually curated by identifying genes present in the CosMx panel, which have been previously implicated as epithelial or mesenchymal related genes (25, 26, 27, 28). GSEA between tumor cell types were performed using the “Hallmark” gene set (31) and the “epithelial” and “mesenchymal” annotated lists with fgsea (81) run on the fold change in expression between groups.

### Cell to cell proximity analyses

Cell to cell proximity analyses were performed measuring the minimum distance in Euclidean distance from the center of each tumor cell to center of the nearest immune cell of question, considering only cells within the same FOV. ANOVA tests were performed with post-hoc (pairwise T-test) of the distance of macrophages or CD8 T-cells to tumor cell types.

### Spatially variable gene (SVG) analysis

PD-1 expression in CD 8 T cells and SPP1 expression in macrophages was examined using spatially variable gene analysis as follows. Negative probes and cells with IDs equal to 0 were removed. Genes with total counts less than 10 or expressed in less than 10% of cells were removed.

Region specific SVG analysis: We implemented SPARK on FOV specific expression matrices along with spatial information to identify FOV specific SVGs. SPARK uses a generalized linear spatial model to assess each gene’s expression pattern, accounting for both spatially structured and unstructured variability. We use the multiplicity corrected adjusted p-values to identify SVGs for each FOV in case p-value < 0.01. Finally, we infer spatial region specific SVGs through intersection of SVGs across FOVs which are annotated for a particular spatial region.

Cell-type specific SVG analysis: To identify spatially varying genes specific to CD8+ or macrophage cells, we employed the SPARK method to a subset of cells which were identified as CD8+ or macrophage across all FOVs. As with the whole-tissue analysis, the significance of spatially varying genes in the CD8+ and macrophage subsets were evaluated using permutation-based p-values and adjusted for multiple comparisons using a threshold of adjusted p-value < 0.01. Results were visualized using “ggplot2”, with spatial maps indicating gene expression variability.

To compare average gene expression in the same cell type across the sarcomatoid, clear cell, and transition regions, we employed the Wilcoxon rank-sum test for pairwise comparisons. The significance threshold was set at p < 0.05.

### GeoMx Spatial Transcriptomics

Specimens 1 – 3 were cut onto slides and regions of interest (ROI’s) were manually selected based on both matched histology from an adjacent H&E, as well as protein markers. Protein markers included PanCK (epithelial cells), CD45 (immune cells), and CD3 (T cells). We segmented regions into immune cells (CD45+) and non-immune cells (CD45-). We analyzed nonimmune cells as tumor cells, recognizing the limitation that there may be some other stromal cells in this compartment, however PanCK was not a reliable stain to isolate tumor cells. The GeoMx whole transcriptome gene panel was used. Gene signatures were applied as detailed below.

GSEA was performed through clusterProfiler(82).

### Multiple Immunofluorescence (mIF) Staining and Analyses

#### mIF staining

mIF was performed as previously described (83). Formalin-fixed, paraffin-embedded (FFPE) tissue slides underwent rehydration three times using xylene, followed by sequential submersion in 100%, 95%, and 70% ethanol, respectively. After rehydration, the slides were washed with neutral buffered formalin and rinsed in deionized water. Antigen retrieval was carried out using sodium citrate buffer (pH 6.0) for membrane and cytoplasmic epitopes and tris-EDTA buffer (pH 9.0) for nuclear epitopes. Tissue was blocked using endogenous peroxidase at room temperature for 1 hr. Primary antibodies, including CD3 (1:100, Abcam), CD8 (1:400, Bioss), CD163 (1:400, Leica), PD-L1 (1:200, Cell Signaling), NCAD (ThermoFisher), and ECAD (Cell Signaling), were diluted in 1% BSA block and incubated for an hour at room temperature. After multiplexing, tissue samples were rinsed with deionized water and mounted using DAPI ProLong™ Diamond Antifade Mountant (Thermo Fisher Scientific). Imaging was performed using the Vectra® Polaris™ Work Station (Akoya Biosciences).

### Quantification of mIF staining

The following multispectral slide scan bands (MOTiF) were used for image capture: DAPI, Opal 480, Opal 520, Opal 570, Opal 620, Opal 690, and Opal 780. After all images were acquired, images were analyzed using inForm® Cell Analysis™ software (Akoya Biosciences). Cell segmentation was performed using DAPI staining to determine cell location and nuclear size, allowing categorization of cells into nucleus and membrane subsets. Automated training software was used to define basic phenotypes including epithelial (N-cadherin low), and mesenchymal (N-cadherin high). The software generated output containing the mean fluorescent intensity (mfi) of each antibody-fluorophore pair, phenotype classifications, and x and y coordinates, were then used for further analysis to assess the relative population of each cell type.

### Effect of N-cadherin expression on cell co-localization

Cell colocalization was quantified for each pair of cell types, using spatial distance metrics calculated using the DIMPLE pipeline (84) from a subset of 342 FOVs across 21 sRCC specimens. We computed the Jensen-Shannon distance (JSD) between each cell’s distributions for each FOV, which are scaled to range from 0 (representing complete separation) to 100 (indicating perfect overlap). N-cadherin expression was also aggregated across cells within FOVs (using mean and total expression). We used linear mixed models to model association between each pairwise cell colocalization (using DIMPLE distances) and N-cadherin (as covariate). To account for the correlation between FOVs within the same scan, we used a random intercept in our model for patient. Additionally, we adjusted for the percentage of each cell type within each FOV and excluded FOVs that didn’t have at least one cell of each type.

### Creating and applying “Clear Cell”, “Transition”, and “Sarcomatoid” Gene signatures

#### Creating gene lists

Differential expression (DE) from CosMx data between Clear Cell and Sarcomatoid was used to create a sarcomatoid gene list and DE between Clear Cell and Transition was used to create a Transition gene list. For both comparisons, DE genes that had an adjusted p-value <0.05 and an absolute log2 fold-change value >0.6 were included in the list.

### Scoring the GeoMx dataset

The GeoMx dataset was normalized by the 3rd quantile (Q3) method. Any rhaboid and outlier ROIs were filtered out before scoring. The ROIs were then scored using the singscore package (85), in which the scores were centered around 0. The scores were then scaled between 0 and 1. The Sarcomatoid gene score was first applied to all histologic sarcomatoid areas, and the lowest Sarcomatoid score in these areas was defined as the Sarcomatoid cutoff. Any clear cell regions with a Sarcomatoid score above the Sarcomatoid cutoff were assigned a transcriptional label of Sarcomatoid. All remaining ROIs were assigned a Transition score. The median Transition score was calculated and any non-Sarcomatoid ROIs at or above this score were labeled as Transition, and any ROIs below this score were labeled as Clear Cell. The same process was performed for each Specimen. GSEA was performed for Clear Cell versus Transition versus Sarcomatoid regions according to their transcriptional singatrue using the Hallmark pathways.

### Scoring and survival analysis for TCGA, CPTAC, and 15G datasets

Before scoring, all datasets were counts per million (CPM )normalized using the edgeR package (86). In TCGA, non-metastatic ccRCC histology samples were used. In CPTAC, metastatic samples were removed. Each dataset was scored using the singscore package (85), which the scores were centered around 0. For survival analysis, we split each dataset to compare the top 25% quantile transition score samples to the rest of the samples. We ran survival analysis on multiple survival time variables (OS, DFI, DSS, PFI). Survival analysis was performed using the survival package (87) and Kaplan-Meier survival plots were created using the ggsurvplot function in the survminer package(88).

### Scoring the Moffitt Dataset

The Moffitt CosMx data was assigned a Transition score at the single cell level for all tumor cells in the same way described above for scoring of the GeoMx dataset, but now at the single cell level. The median Transition score for each region was compared between clear cell regions from pure ccRCC tumors and clear cell regions from sarcomatoid tumors by simple t-test of the means. M2 macrophage content in each region was calculated as a percentage of all cells, and regression performed to correlated macrophage percentage with median Transition score incorporating both the clear cell regions from pure ccRCC tumors and from sarcomatoid tumors.

### Cell culture

786-O and THP-1 were cultured in RPMI medium containing 10% FBS and penicillin– streptomycin (15140-122, Invitrogen). UOK127 was cultured in DMEM medium containing 10% FBS and P/S supplemented with non-essential amino acids (11140050, Thermo Fisher Scientific). Cells were cultured at 37°C in 5% CO2. 786-O and THP-1 were obtained from American Type Culture Collection (ATCC), and UOK127 cells were obtained from Dr. Marston Linehan (National Institutes of Health).

### Antibodies and chemicals for in vitro experiments

Primary antibodies for E-cadherin (3195), Snail (3879), Slug (9585), CD44 (3570), Met (8198), ANXA2 (8235), PD-L1 (29122), and GAPDH (5174) were purchased from Cell Signaling Technology, and primary antibodies for CD163 (ab182422), CD206 (ab64693), and SPP1 (ab69498) were purchased from Abcam. Primary antibodies for N-cadherin (610920), CCL20 (PA5-95917), and α-tubulin (T6074) were purchased from Becton Dickinson, Invitrogen, and Sigma-Aldrich, respectively. HRP conjugated secondary antibodies, anti-rabbit (711-035-152) and anti-mouse (115-035-146), were purchased from Jackson Immunoresearch. We purchased Recombinant Human (rh) TGF-beta 1 (7754-BH), rh SPP1 (1433-OP), and rh CCL20 (360-MP) from R&D systems, rh Interleukin-4 (IL-4) (ab155733) from Abcam, lipopolysaccharide (LPS) (L4391) from Sigma-Aldrich, and rh Interferon (IFN)-γ (PHC4031) from Thermo Fisher Scientific.

### Preparation of macrophage-like cells

Macrophage-like cells were generated from THP-1, a human monocytic leukemia cell line. THP-1 cells (1.0 × 10^6^ cells/mL for 100mm dish or 3.0 × 10^5^ cells/mL for 6 well cell culture inserts) were treated with 100 ng/mL phorbol 12-myristate 13-acetate (PMA, P8139, Sigma-Aldrich) for 24 h. After the cells attached to the plate, they were washed three times with PBS to remove PMA and cultured in regular RPMI medium for 1 day then transferred to new plates.

Macrophages cultured in regular medium for another 2 days were defined as M0, those cultured with 10 ng/mL LPS and 20 ng/mL rh IFN-γ were defined as M1, and those cultured with 20 ng/mL rhIL-4 were defined as M2.

### Macrophage-conditioned medium (CM)

The macrophage-like cells were cultured in RPMI medium containing 10% FBS for 48 hours. The collected medium was then centrifuged at 1500 g for 10 minutes, and the supernatant was used as CM.

### Invasion assays

Invasion assay kits using Matrigel (354480, Corning) were used for invasion assays in 24-well culture plates. The upper chamber contained (1.0 × 10 ^4^ 786-O or 2.0 × 10 ^4^ UOK127) cells in 300 μL of serum-free medium, and the lower chamber was filled with 700 μL of medium containing 10% FBS with or without M2 CM. In the lower chamber containing M2 CM, CM accounted for 30% of the 700 μL of medium.

The cells were incubated at 37°C and 5% CO2 for 24 hours. Cells present on the upper surface of the cell culture insert were removed with a cotton swab, and cells on the lower surface were stained with Diff-Quik. Invading cells were counted manually in five randomly selected 200x fields. Values are presented as mean ± SEM. Statistical analysis was performed using Student’s t-test to determine significant differences between groups. Statistical analysis was performed using Prism 10 (GraphPad).

### Western blotting

Cell lysates were prepared using RIPA buffer (89900, Thermo Fisher Scientific) containing protease and phosphatase inhibitor cocktail (P0044, P7626 and P8340, Sigma-Aldrich). Lysates were separated by SDS-PAGE, transferred to nitrocellulose or PVDF membranes and blocked for 1 h at room temperature in 0.05% Tween Tris-buffered saline (TBST) with 3% nonfat dry milk (1706404, Bio-Rad). Membranes were incubated overnight at 4°C with primary antibodies diluted in 3% BSA in TBST. After washing three times, membranes were incubated with HRP-conjugated anti-rabbit or anti-mouse secondary antibodies for 1 h at room temperature. After washing three times again, protein bands were detected using ECL Substrate (1705062, Bio-Rad).

### siRNA transfection

For transfection, lipofection was performed using jetPRIME® (101000015, Polyplus). Lipofection was performed on macrophage-like cells seeded at a density of 3 × 10^5^ cells/well on 6-well cell culture inserts. They were incubated for 48 hours at 50 nM siRNA according to the protocol provided by the vendor.

Negative control siRNA (4390844), SPP1 siRNA #1 (4392420, s13375) and SPP1 siRNA #2 (4392420, s13376) were purchased from Thermo Fisher Scientific.

## Resource availability

### Data availability

Raw data from CosMx and GeoMx analyses are deposited in Gene Expression Omnibus (GEO) as [GSE299368 and GSE299598 respectively] and will be publicly available as of the date of publication.

Original western blot images will be deposited at Mendeley and publicly available as of the date of publication.

Multiplex immunofluorescent, H&E, and microscopy data will be shared by the lead contact upon request.

Any additional information required to re-analyze the data in this paper is available from the lead contact upon request.

### Code availability

All original code is deposited at Zenodo and publicly available (doi 10.5281/zenodo.15643039).

*Lead contact:* (E.T.K)

## Acknowledgements

This study was supported by funding through the Forbes Institute for Discovery at University of Michigan, the Rogel Cancer Center: TrEC Scholarship Graduate Student Scholarship (to Jessica Aldous) and through the Clark Family Fellowship for Kidney Cancer Research (to Allison May). Research reported in this publication was supported by a National Institute of Health Training Grant T32F054350 (to Allison May) and the National Cancer Institute under Award Number P30 CA046592 by the use of the following Cancer Center Shared Resource(s):Single Cell Spatial Analysis Shared Resource, Tissue Molecular Pathology Shared Resource and Cancer Data Science Shared Resource. UOK-127 cells were gifted by the lab of Dr. Marston Linehan at the National Cancer Institute.

## Disclosure of Conflict of Interest

CW is an employee of Bruker Spatial Biology and hold stock and/or stock options in the company.

## SUPPLEMENTAL

**Figure S1.**
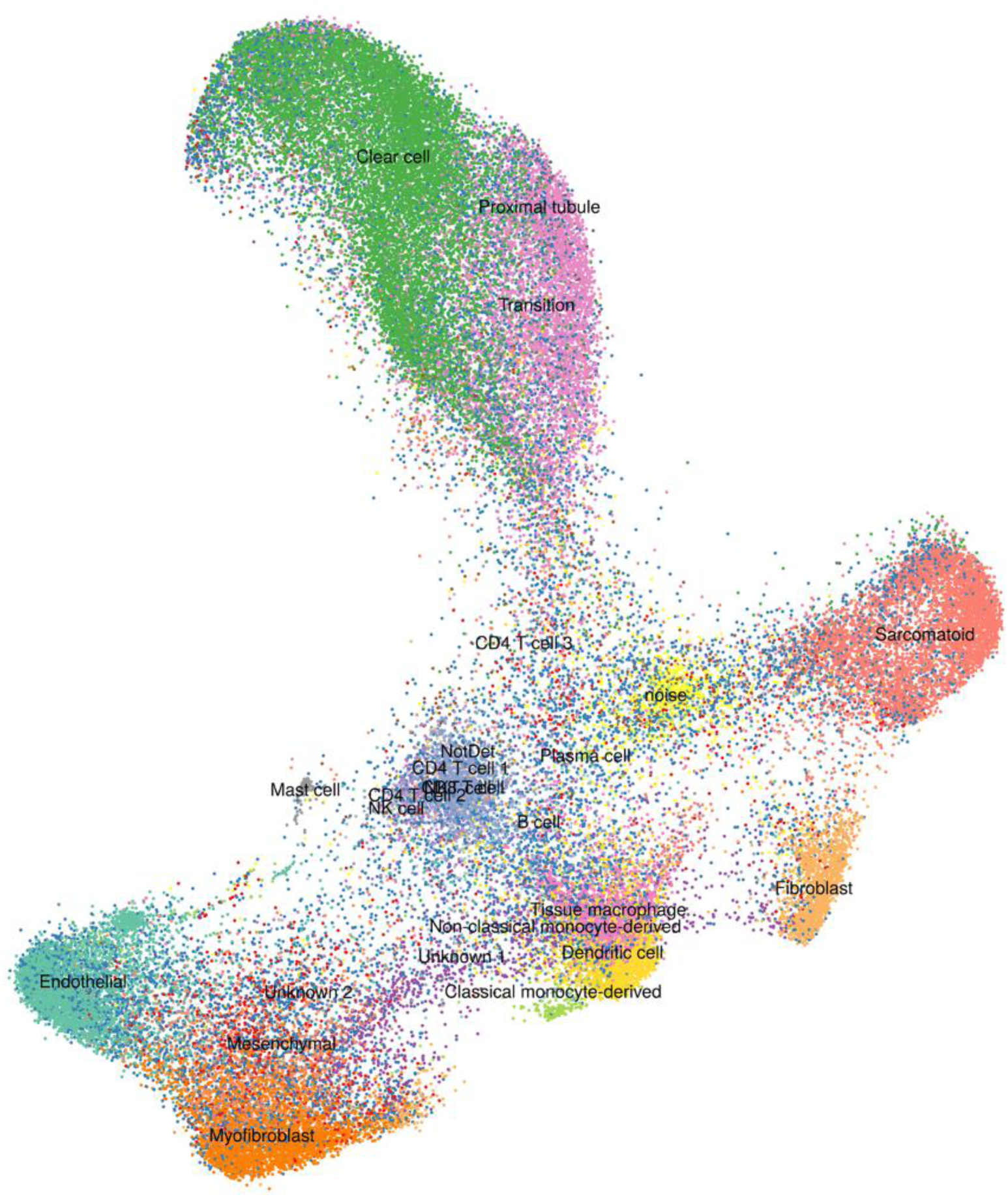
High definition UMAP of malignant and non-malignant cells captured across all FOVs from CosMx analysis of Specimen 1.

**Figure S2.**
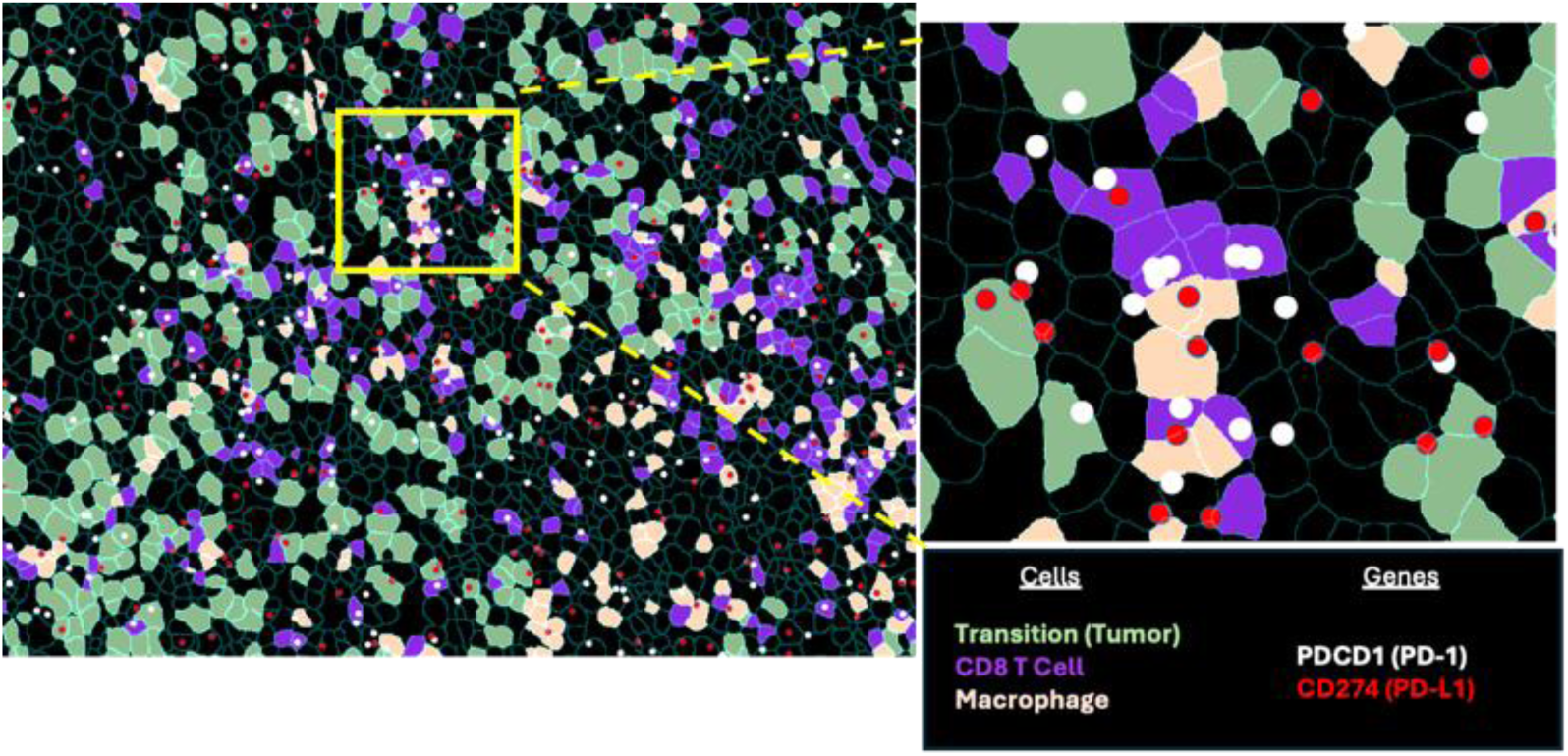
Representative field of view from CosMx analysis showing cell types based on transcriptional cluster and gene expression of PD-1 and PD-L1.

**Figure S3.**
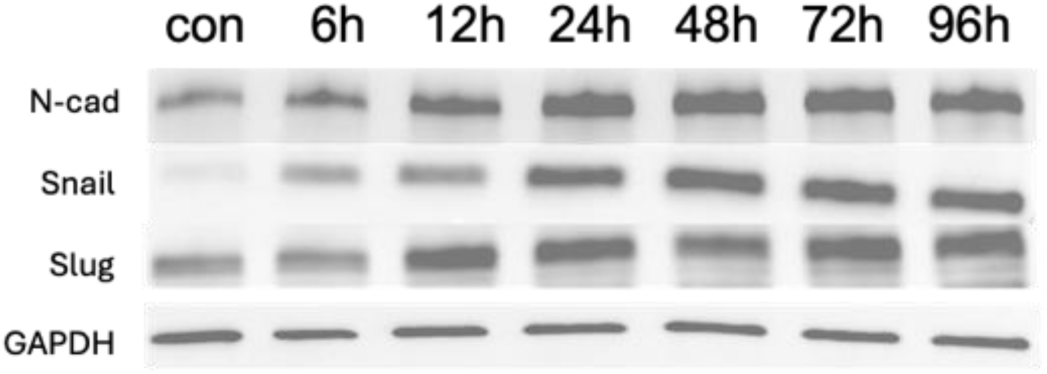
Western blot showing EMT markers in 786-O cells treated with 20ng/mL TGFB at varying time points.

**Figure S4.**
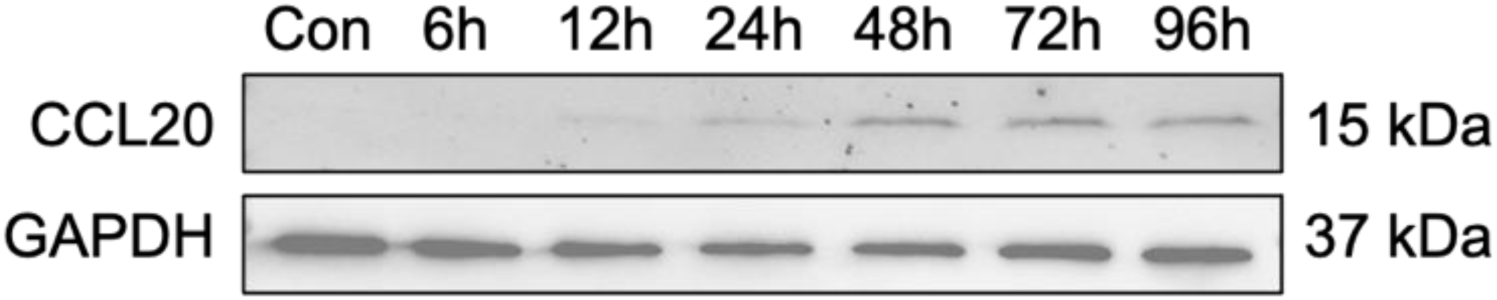
Western blot showing CCL20 expression in UOK-127 cells treated with 20ng/mL recombinant TGFB at varying timepoints.

**Figure S5.**
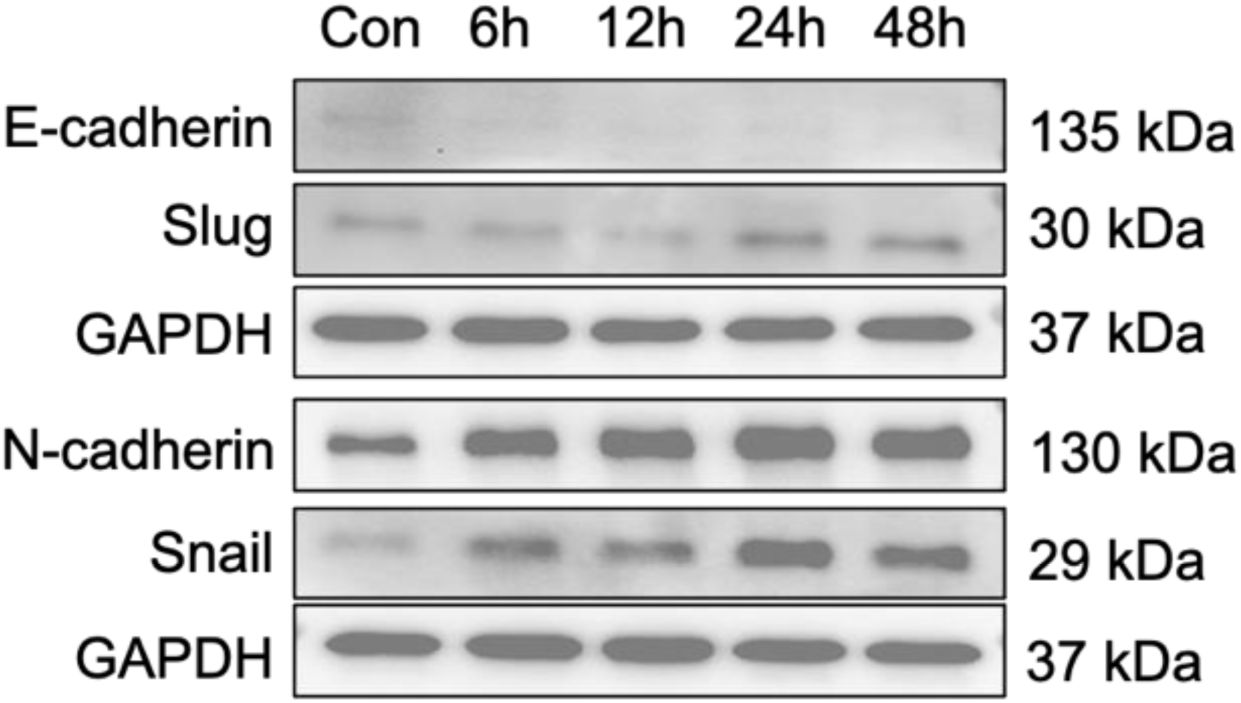
Western blot showing EMT markers in UOK-127 cells treated with 50ng/mL recombinant SPP1 at varying timepoints.

**Figure S6.**
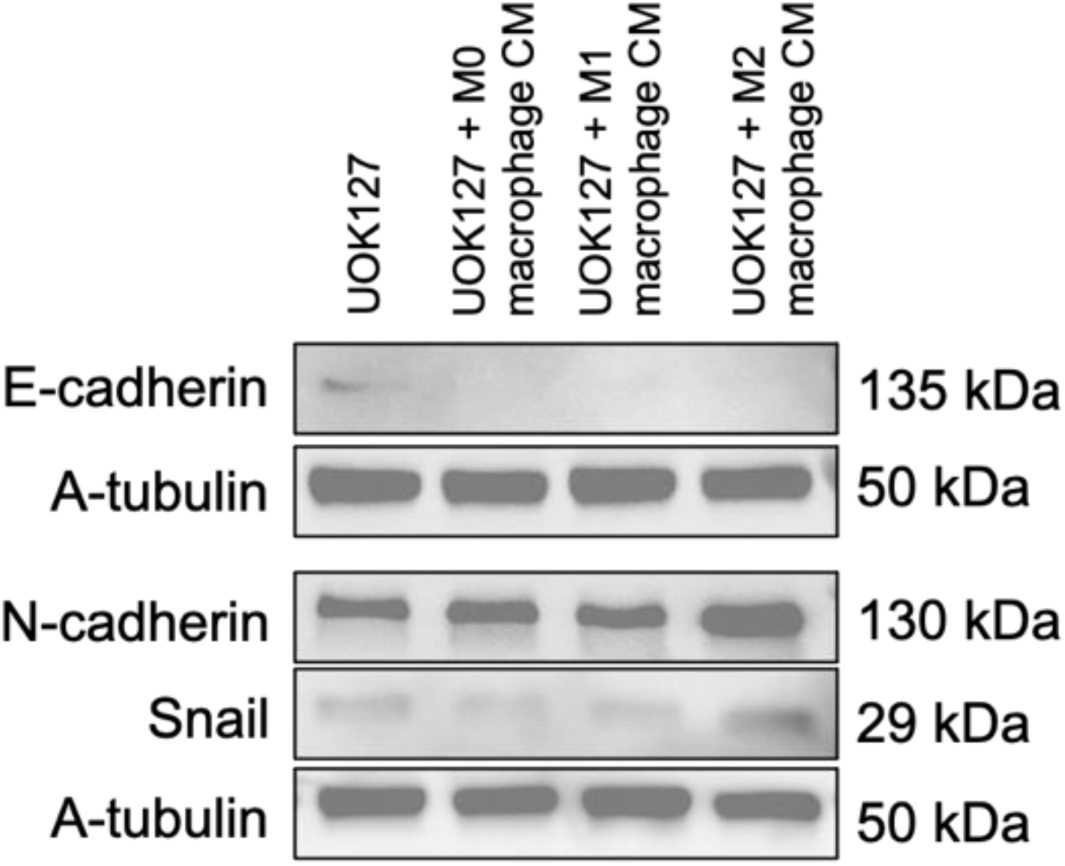
Western blot showing EMT markers in UOK-127 cells treated with macrophage conditioned media (CM).

**Figure S7.**
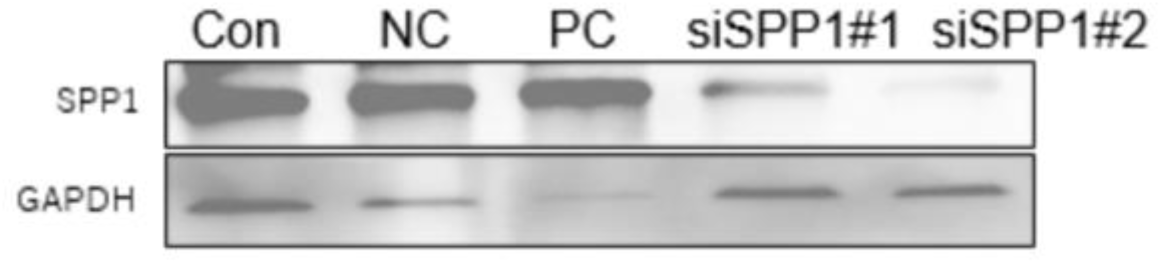
Western blot showing SPP1 expression in THP-1 derived M2-like macrophages including control, and treatment with silencing mRNAs siNC, siPC, siSPP1#1, and siSPP1#2.

**Figure S8.**
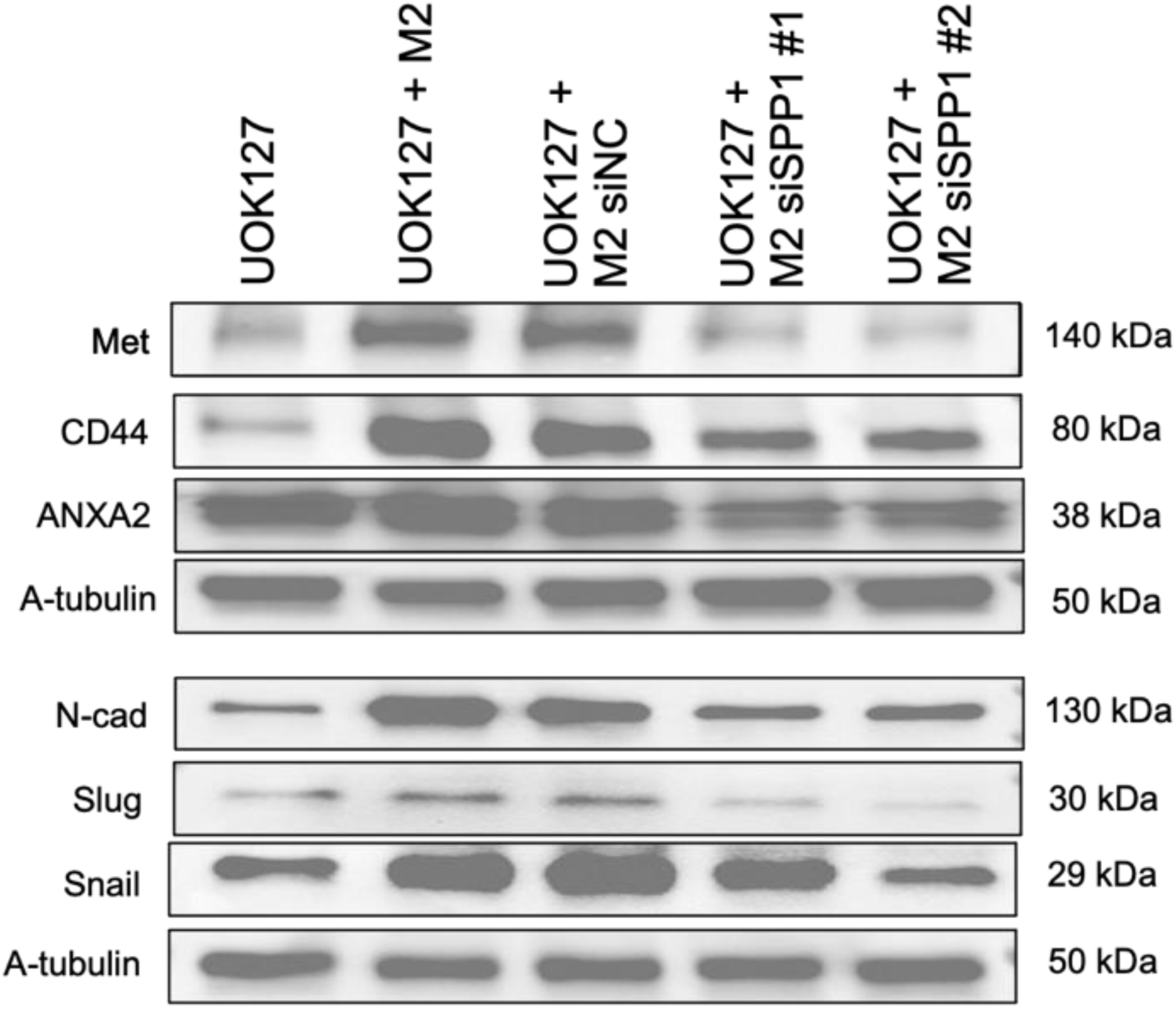
Western blot showing sarcomatoid-specific proteins (Met, CD44, ANXA2) and EMT markers (N-cadherin, Slug, Snail) in UOK-127 cells control, treated with M2-like macrophages, or treated with M2-like macrophages with siRNA knockdown of negative control or SPP1.

**Figure S9.**
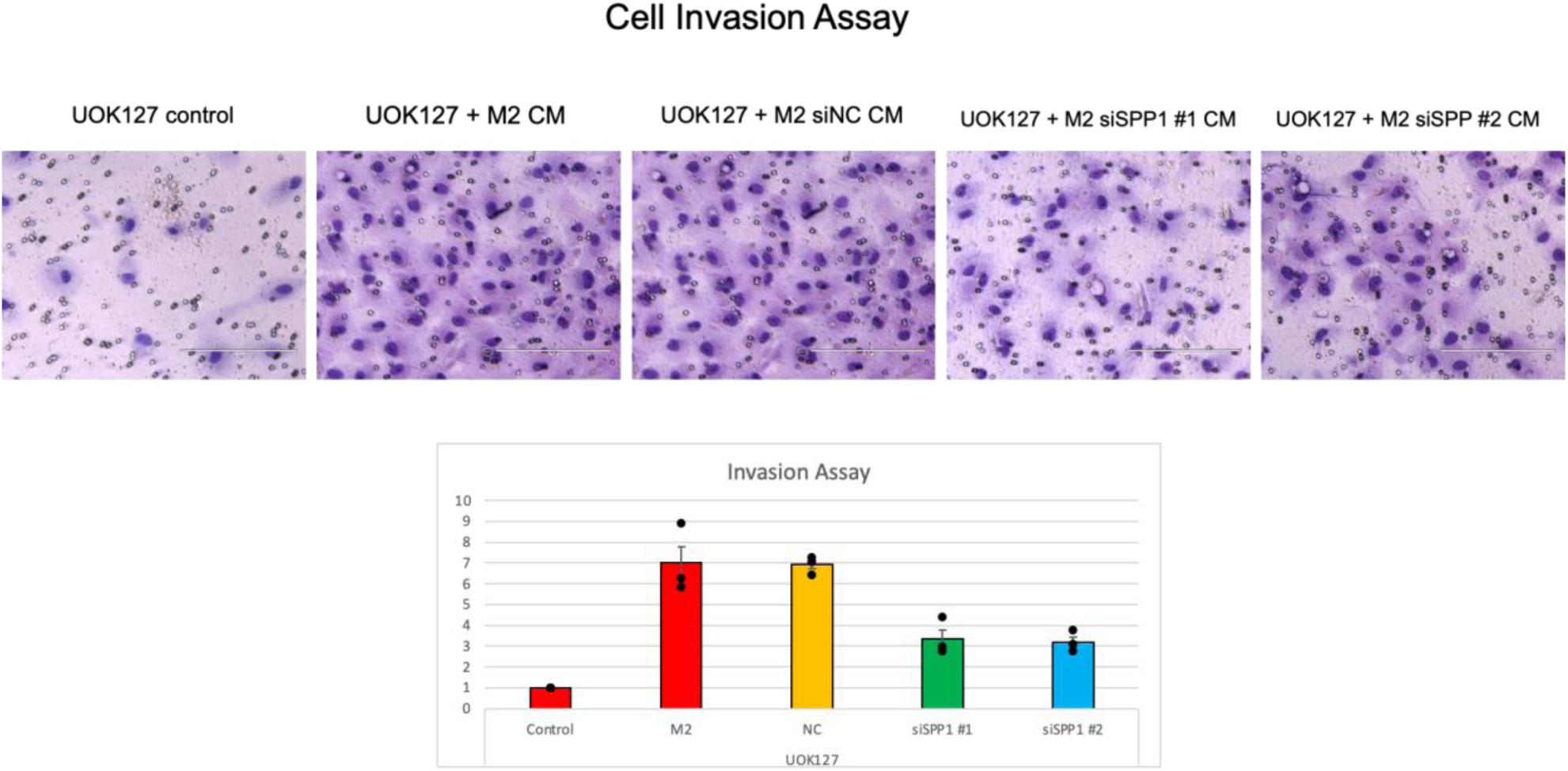
Invasion assay of UOK-127 cell co-cultured with M2 macrophage conditioned media with negative controle (siNC) or silencing of SPP1 (siSPP1) in M2 macrophages.

**Table S1.**
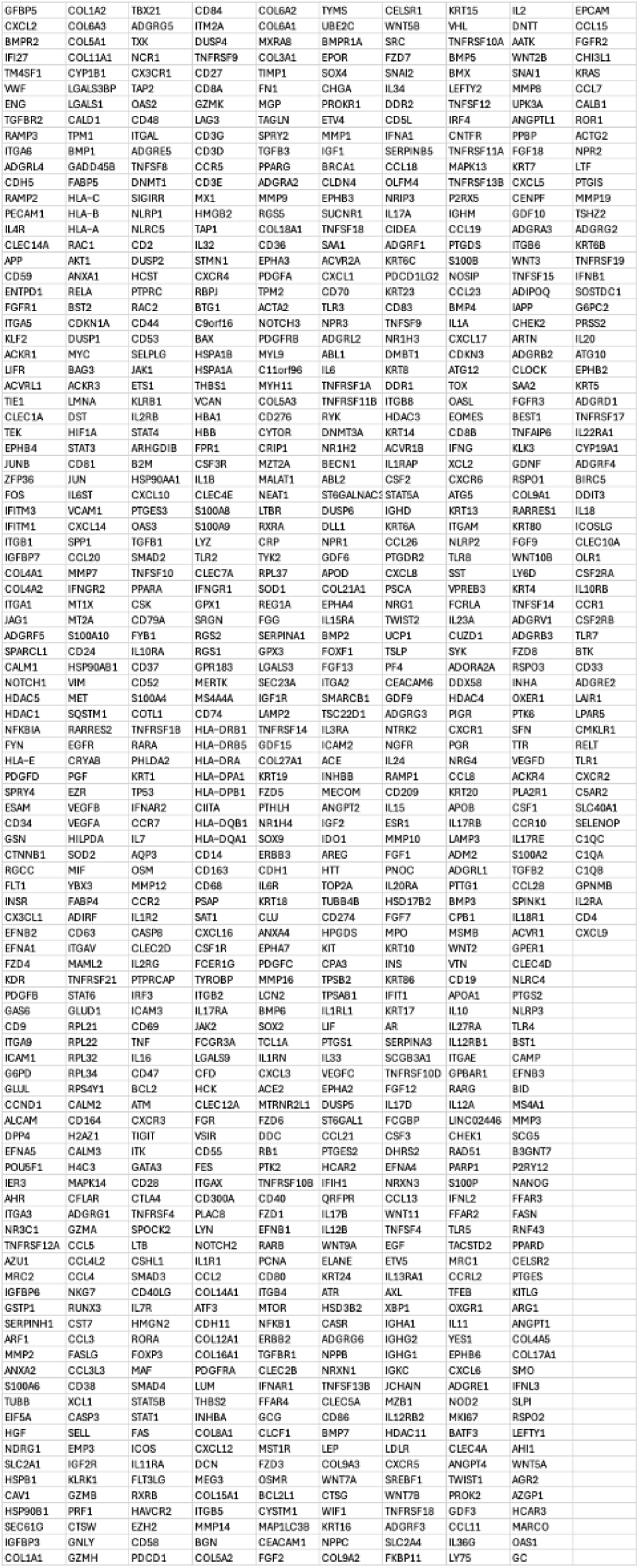
List of genes in 960-gene panel from CosMx analysis.

**Table S2.** List of manually curated “epithelial” and “mesenchymal” gene list from CosMx gene panel.

| Epithelial | Mesenchymal |
| --- | --- |
| AGR2 | AXL |
| AQP3 | BGN |
| ARHGDIB | C9orf19 |
| AZGP1 | CALD1 |
| CD24 | CAV1 |
| CDH1 | CCL2 |
| CEACAM1 | CCL8 |
| CEACAM6 | CD163 |
| COL17A1 | CDH11 |
| DDR1 | CLEC2B |
| EPCAM | COL14A1 |
| ERBB2 | COL15A1 |
| ERBB3 | COL1A1 |
| EZR | COL1A2 |
| FGFR3 | COL3A1 |
| GDF15 | COL5A2 |
| GPR110 | COL6A1 |
| IL1B | COL6A2 |
| IL20RA | CRYAB |
| IL4R | CSF2RB |
| ITGB6 | CXCL12 |
| KRT14 | CXCR4 |
| KRT15 | CYP1B1 |
| KRT17 | DCN |
| KRT18 | DDR2 |
| KRT19 | EMP3 |
| KRT5 | FN1 |
| KRT6A | FYN |
| KRT6B | GZMK |
| KRT7 | IGF1 |
| KRT8 | IGFBP3 |
| MAPK13 | IGFBP5 |
| MPP7 | IL10RA |
| MST1R | ITM2A |
| PHLDA2 | MAF |
| PSCA | MMP2 |
| PTK6 | MRC1 |
| S100A8 | MYL9 |
| S100P | NR3C1 |
| SAA1 | PDGFC |
| SLPI | PPARG |
| TACSTD2 | PTGDS |
|  | PTGIS |
|  | PTPRC |
|  | RARRES2 |
|  | RGS2 |
|  | ROR1 |
|  | SNAI2 |
|  | SPARCL1 |
|  | SRGN |
|  | TAGLN |
|  | TGFBI |
|  | TNFAIP6 |
|  | TNFRSF21 |
|  | TPM2 |
|  | TWIST1 |
|  | VCAM1 |
|  | VCAN |
|  | VIM |
|  | WNT5A |

**Table S3.**
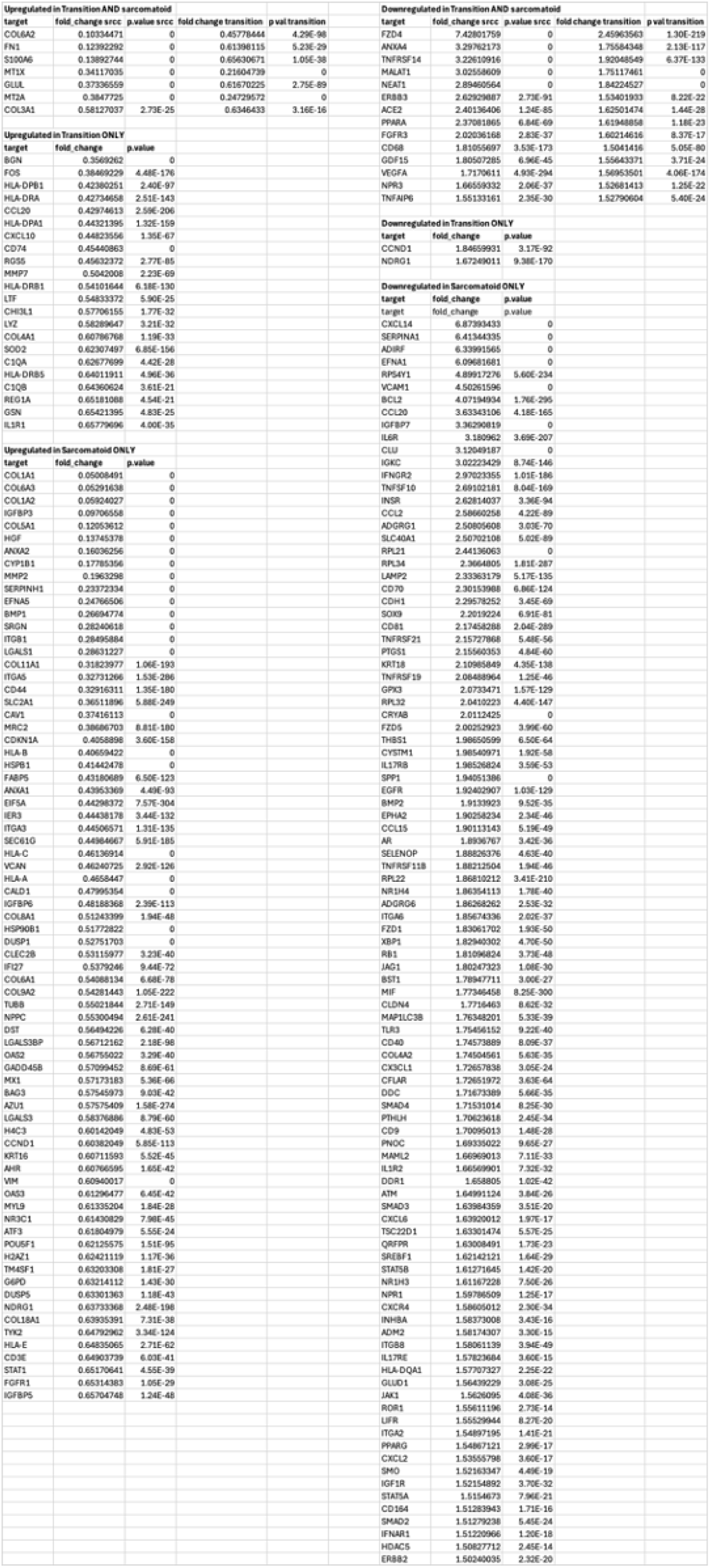
All genes up or downregulated in differential gene expression between Clear Cells and Transition cells and Clear Cells and Sarcomatoid cells in CosMx analysis, including fold change and p-value.

**Table S4.**
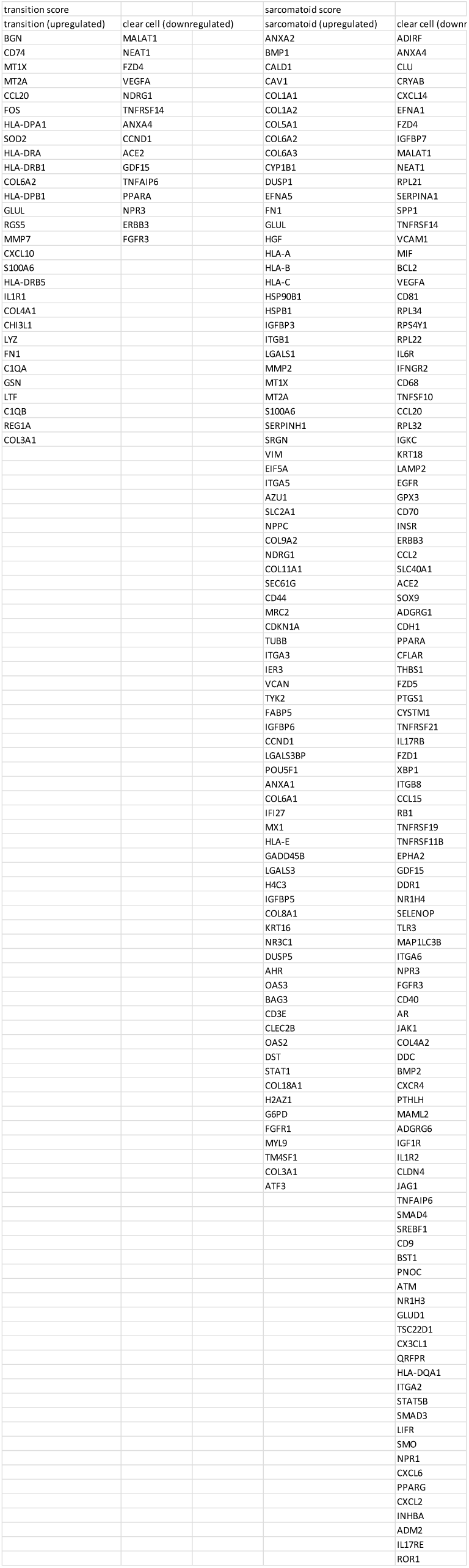
Genes comprising Transition and Sarcomatoid gene signatures.

## Literature cited

1. Siegel RL, Kratzer TB, Giaquinto AN, Sung H, Jemal A. Cancer statistics, 2025. CA: A Cancer Journal for Clinicians. 2025;75(1):10-45.

2. Rao A, Wiggins C, Lauer RC. Survival outcomes for advanced kidney cancer patients in the era of targeted therapies. Ann Transl Med. 2018;6(9):165.

3. Siegel RL, Miller KD, Fuchs HE, Jemal A. Cancer statistics, 2022. CA Cancer J Clin. 2022;72(1):7-33.

4. Blum KA, Gupta S, Tickoo SK, Chan TA, Russo P, Motzer RJ, et al. Sarcomatoid renal cell carcinoma: biology, natural history and management. Nat Rev Urol. 2020;17(12):659–78.

5. Abel EJ, Carrasco A, Culp SH, Matin SF, Tamboli P, Tannir NM, et al. Limitations of preoperative biopsy in patients with metastatic renal cell carcinoma: comparison to surgical pathology in 405 cases. BJU Int. 2012;110(11):1742–6.

6. Mikami S, Katsube K, Oya M, Ishida M, Kosaka T, Mizuno R, et al. Expression of Snail and Slug in renal cell carcinoma: E-cadherin repressor Snail is associated with cancer invasion and prognosis. Lab Invest. 2011;91(10):1443–58.

7. Conant JL, Peng Z, Evans MF, Naud S, Cooper K. Sarcomatoid renal cell carcinoma is an example of epithelial--mesenchymal transition. J Clin Pathol. 2011;64(12):1088–92.

8. Taki M, Abiko K, Ukita M, Murakami R, Yamanoi K, Yamaguchi K, et al. Tumor Immune Microenvironment during Epithelial-Mesenchymal Transition. Clin Cancer Res. 2021;27(17):4669–79.

9. Greaves D, Calle Y. Epithelial Mesenchymal Transition (EMT) and Associated Invasive Adhesions in Solid and Haematological Tumours. Cells. 2022;11(4).

10. Bi M, Zhao S, Said JW, Merino MJ, Adeniran AJ, Xie Z, et al. Genomic characterization of sarcomatoid transformation in clear cell renal cell carcinoma. Proc Natl Acad Sci U S A. 2016;113(8):2170–5.

11. Malouf GG, Ali SM, Wang K, Balasubramanian S, Ross JS, Miller VA, et al. Genomic Characterization of Renal Cell Carcinoma with Sarcomatoid Dedifferentiation Pinpoints Recurrent Genomic Alterations. Eur Urol. 2016;70(2):348–57.

12. Wang Z, Kim TB, Peng B, Karam J, Creighton C, Joon A, et al. Sarcomatoid Renal Cell Carcinoma Has a Distinct Molecular Pathogenesis, Driver Mutation Profile, and Transcriptional Landscape. Clin Cancer Res. 2017;23(21):6686–96.

13. Bakouny Z, Braun DA, Shukla SA, Pan W, Gao X, Hou Y, et al. Integrative molecular characterization of sarcomatoid and rhabdoid renal cell carcinoma. Nat Commun. 2021;12(1):808.

14. Tickoo SK, Alden D, Olgac S, Fine SW, Russo P, Kondagunta GV, et al. Immunohistochemical expression of hypoxia inducible factor-1alpha and its downstream molecules in sarcomatoid renal cell carcinoma. J Urol. 2007;177(4):1258–63.

15. Tannir NM, Signoretti S, Choueiri TK, McDermott DF, Motzer RJ, Flaifel A, et al. Efficacy and Safety of Nivolumab Plus Ipilimumab versus Sunitinib in First-line Treatment of Patients with Advanced Sarcomatoid Renal Cell Carcinoma. Clin Cancer Res. 2021;27(1):78–86.

16. Rini BI, Motzer RJ, Powles T, McDermott DF, Escudier B, Donskov F, et al. Atezolizumab plus Bevacizumab Versus Sunitinib for Patients with Untreated Metastatic Renal Cell Carcinoma and Sarcomatoid Features: A Prespecified Subgroup Analysis of the IMmotion151 Clinical Trial. Eur Urol. 2021;79(5):659–62.

17. Wang G, Xu D, Zhang Z, Li X, Shi J, Sun J, et al. The pan-cancer landscape of crosstalk between epithelial-mesenchymal transition and immune evasion relevant to prognosis and immunotherapy response. NPJ Precis Oncol. 2021;5(1):56.

18. Romeo E, Caserta CA, Rumio C, Marcucci F. The Vicious Cross-Talk between Tumor Cells with an EMT Phenotype and Cells of the Immune System. Cells. 2019;8(5).

19. Raskov H, Orhan A, Christensen JP, Gogenur I. Cytotoxic CD8(+) T cells in cancer and cancer immunotherapy. Br J Cancer. 2021;124(2):359–67.

20. Choueiri TK, Tomczak P, Park SH, Venugopal B, Ferguson T, Symeonides SN, et al. Overall Survival with Adjuvant Pembrolizumab in Renal-Cell Carcinoma. N Engl J Med. 2024;390(15):1359–71.

21. Rosellini M, Marchetti A, Mollica V, Rizzo A, Santoni M, Massari F. Prognostic and predictive biomarkers for immunotherapy in advanced renal cell carcinoma. Nat Rev Urol. 2023;20(3):133–57.

22. Mandal M, Ghosh B, Anura A, Mitra P, Pathak T, Chatterjee J. Modeling continuum of epithelial mesenchymal transition plasticity. Integr Biol (Camb). 2016;8(2):167–76.

23. Simeonov KP, Byrns CN, Clark ML, Norgard RJ, Martin B, Stanger BZ, et al. Single-cell lineage tracing of metastatic cancer reveals selection of hybrid EMT states. Cancer Cell. 2021;39(8):1150–62 e9.

24. He S, Bhatt R, Brown C, Brown EA, Buhr DL, Chantranuvatana K, et al. High-plex imaging of RNA and proteins at subcellular resolution in fixed tissue by spatial molecular imaging. Nat Biotechnol. 2022;40(12):1794–806.

25. Mlcochova H, Machackova T, Rabien A, Radova L, Fabian P, Iliev R, et al. Epithelial-mesenchymal transition-associated microRNA/mRNA signature is linked to metastasis and prognosis in clear-cell renal cell carcinoma. Sci Rep. 2016;6:31852.

26. Byers LA, Diao L, Wang J, Saintigny P, Girard L, Peyton M, et al. An epithelial-mesenchymal transition gene signature predicts resistance to EGFR and PI3K inhibitors and identifies Axl as a therapeutic target for overcoming EGFR inhibitor resistance. Clin Cancer Res. 2013;19(1):279–90.

27. Tan TZ, Miow QH, Miki Y, Noda T, Mori S, Huang RY, et al. Epithelial-mesenchymal transition spectrum quantification and its efficacy in deciphering survival and drug responses of cancer patients. EMBO Mol Med. 2014;6(10):1279–93.

28. George JT, Jolly MK, Xu S, Somarelli JA, Levine H. Survival Outcomes in Cancer Patients Predicted by a Partial EMT Gene Expression Scoring Metric. Cancer Res. 2017;77(22):6415–28.

29. Delahunt B, Bethwaite PB, McCredie MR, Nacey JN. The evolution of collagen expression in sarcomatoid renal cell carcinoma. Hum Pathol. 2007;38(9):1372–7.

30. Subramanian A, Tamayo P, Mootha VK, Mukherjee S, Ebert BL, Gillette MA, et al. Gene set enrichment analysis: a knowledge-based approach for interpreting genome-wide expression profiles. Proc Natl Acad Sci U S A. 2005;102(43):15545–50.

31. Liberzon A, Birger C, Thorvaldsdottir H, Ghandi M, Mesirov JP, Tamayo P. The Molecular Signatures Database (MSigDB) hallmark gene set collection. Cell Syst. 2015;1(6):417–25.

32. Motzer RJ, Banchereau R, Hamidi H, Powles T, McDermott D, Atkins MB, et al. Molecular Subsets in Renal Cancer Determine Outcome to Checkpoint and Angiogenesis Blockade. Cancer Cell. 2020;38(6):803–17 e4.

33. Zhu J, Lin Q, Zheng H, Rao Y, Ji T. The pro-invasive factor COL6A2 serves as a novel prognostic marker of glioma. Front Oncol. 2022;12:897042.

34. Li B, Shen W, Peng H, Li Y, Chen F, Zheng L, et al. Fibronectin 1 promotes melanoma proliferation and metastasis by inhibiting apoptosis and regulating EMT. Onco Targets Ther. 2019;12:3207–21.

35. Zhang C, Zeng M, Xu Y, Huang B, Shi P, Zhu X, et al. S100A6 mediated epithelial-mesenchymal transition affects chemosensitivity of colorectal cancer to oxaliplatin. Gene. 2024;914:148406.

36. Lei Y, Yan W, Lin Z, Liu J, Tian D, Han P. Comprehensive analysis of partial epithelial mesenchymal transition-related genes in hepatocellular carcinoma. J Cell Mol Med. 2021;25(1):448–62.

37. Paul I, Bolzan D, Youssef A, Gagnon KA, Hook H, Karemore G, et al. Parallelized multidimensional analytic framework applied to mammary epithelial cells uncovers regulatory principles in EMT. Nat Commun. 2023;14(1):688.

38. Lafitte M, Moranvillier I, Garcia S, Peuchant E, Iovanna J, Rousseau B, et al. FGFR3 has tumor suppressor properties in cells with epithelial phenotype. Mol Cancer. 2013;12:83.

39. Yao H, Sun C, Hu Z, Wang W. The role of annexin A4 in cancer. Front Biosci (Landmark Ed). 2016;21(5):949–57.

40. Liu Q, Song M, Wang Y, Zhang P, Zhang H. CCL20-CCR6 signaling in tumor microenvironment: Functional roles, mechanisms, and immunotherapy targeting. Biochim Biophys Acta Rev Cancer. 2025;1880(3):189341.

41. Vazirinejad R, Ahmadi Z, Kazemi Arababadi M, Hassanshahi G, Kennedy D. The biological functions, structure and sources of CXCL10 and its outstanding part in the pathophysiology of multiple sclerosis. Neuroimmunomodulation. 2014;21(6):322–30.

42. Zhang W, Borcherding N, Kolb R. IL-1 Signaling in Tumor Microenvironment. Adv Exp Med Biol. 2020;1240:1–23.

43. Yu JE, Yeo IJ, Han SB, Yun J, Kim B, Yong YJ, et al. Significance of chitinase-3-like protein 1 in the pathogenesis of inflammatory diseases and cancer. Exp Mol Med. 2024;56(1):1–18.

44. Ma B, Akosman B, Kamle S, Lee CM, He CH, Koo JS, et al. CHI3L1 regulates PD-L1 and anti-CHI3L1-PD-1 antibody elicits synergistic antitumor responses. J Clin Invest. 2021;131(21).

45. Alspach E, Lussier DM, Miceli AP, Kizhvatov I, DuPage M, Luoma AM, et al. MHC-II neoantigens shape tumour immunity and response to immunotherapy. Nature. 2019;574(7780):696-701.

46. Wu X, Li T, Jiang R, Yang X, Guo H, Yang R. Targeting MHC-I molecules for cancer: function, mechanism, and therapeutic prospects. Mol Cancer. 2023;22(1):194.

47. Maeda M, Johnson KR, Wheelock MJ. Cadherin switching: essential for behavioral but not morphological changes during an epithelium-to-mesenchyme transition. J Cell Sci. 2005;118(Pt 5):873–87.

48. Hollestelle A, Peeters JK, Smid M, Timmermans M, Verhoog LC, Westenend PJ, et al. Loss of E-cadherin is not a necessity for epithelial to mesenchymal transition in human breast cancer. Breast Cancer Res Treat. 2013;138(1):47–57.

49. Kawakami F, Sircar K, Rodriguez-Canales J, Fellman BM, Urbauer DL, Tamboli P, et al. Programmed cell death ligand 1 and tumor-infiltrating lymphocyte status in patients with renal cell carcinoma and sarcomatoid dedifferentiation. Cancer. 2017;123(24):4823–31.

50. Jiang Y, Zhan H. Communication between EMT and PD-L1 signaling: New insights into tumor immune evasion. Cancer Lett. 2020;468:72–81.

51. Soupir AC, Hayes MT, Peak TC, Ospina O, Chakiryan NH, Berglund AE, et al. Increased spatial coupling of integrin and collagen IV in the immunoresistant clear-cell renal-cell carcinoma tumor microenvironment. Genome Biol. 2024;25(1):308.

52. Ricketts CJ, De Cubas AA, Fan H, Smith CC, Lang M, Reznik E, et al. The Cancer Genome Atlas Comprehensive Molecular Characterization of Renal Cell Carcinoma. Cell Rep. 2018;23(1):313–26 e5.

53. Clark DJ, Dhanasekaran SM, Petralia F, Pan J, Song X, Hu Y, et al. Integrated Proteogenomic Characterization of Clear Cell Renal Cell Carcinoma. Cell. 2019;179(4):964–83 e31.

54. Mehra R, Nallandhighal S, Cotta B, Knuth Z, Su F, Kasputis A, et al. Discovery and Validation of a 15-Gene Prognostic Signature for Clear Cell Renal Cell Carcinoma. JCO Precis Oncol. 2024;8:e2300565.

55. Shigeoka M, Koma YI, Kodama T, Nishio M, Akashi M, Yokozaki H. Tongue Cancer Cell-Derived CCL20 Induced by Interaction With Macrophages Promotes CD163 Expression on Macrophages. Front Oncol. 2021;11:667174.

56. Zhang R, Dong M, Tu J, Li F, Deng Q, Xu J, et al. PMN-MDSCs modulated by CCL20 from cancer cells promoted breast cancer cell stemness through CXCL2-CXCR2 pathway. Signal Transduct Target Ther. 2023;8(1):97.

57. Yang Z, Xie H, He D, Li L. Infiltrating macrophages increase RCC epithelial mesenchymal transition (EMT) and stem cell-like populations via AKT and mTOR signaling. Oncotarget. 2016;7(28):44478–91.

58. Xu C, Sun L, Jiang C, Zhou H, Gu L, Liu Y, et al. SPP1, analyzed by bioinformatics methods, promotes the metastasis in colorectal cancer by activating EMT pathway. Biomed Pharmacother. 2017;91:1167–77.

59. Yang Y, Gong Y, Ding Y, Sun S, Bai R, Zhuo S, et al. LINC01133 promotes pancreatic ductal adenocarcinoma epithelial-mesenchymal transition mediated by SPP1 through binding to Arp3. Cell Death Dis. 2024;15(7):492.

60. Ding Y, Fang J, Chen M, Xu Y, Liu N, Fang S, et al. MT1X is an oncogene and indicates prognosis in ccRCC. Biosci Rep. 2022;42(10).

61. Wang J, Zuo Z, Yu Z, Chen Z, Meng X, Ma Z, et al. Single-cell transcriptome analysis revealing the intratumoral heterogeneity of ccRCC and validation of MT2A in pathogenesis. Funct Integr Genomics. 2023;23(4):300.

62. Ma J, Wu R, Chen Z, Zhang Y, Zhai W, Zhu R, et al. CD44 Is a Prognostic Biomarker Correlated With Immune Infiltrates and Metastasis in Clear Cell Renal Cell Carcinoma. Anticancer Res. 2023;43(8):3493–506.

63. Hsieh CH, Hsiung SC, Yeh CT, Yen CF, Chou YW, Lei WY, et al. Differential expression of CD44 and CD24 markers discriminates the epitheliod from the fibroblastoid subset in a sarcomatoid renal carcinoma cell line: evidence suggesting the existence of cancer stem cells in both subsets as studied with sorted cells. Oncotarget. 2017;8(9):15593–609.

64. Yang SF, Hsu HL, Chao TK, Hsiao CJ, Lin YF, Cheng CW. Annexin A2 in renal cell carcinoma: expression, function, and prognostic significance. Urol Oncol. 2015;33(1):22 e11-22 e1.

65. El Zarif T, Semaan K, Eid M, Seo JH, Garinet S, Davidsohn MP, et al. Epigenomic signatures of sarcomatoid differentiation to guide the treatment of renal cell carcinoma. Cell Rep. 2024;43(6):114350.

66. Joseph RW, Millis SZ, Carballido EM, Bryant D, Gatalica Z, Reddy S, et al. PD-1 and PD-L1 Expression in Renal Cell Carcinoma with Sarcomatoid Differentiation. Cancer Immunol Res. 2015;3(12):1303–7.

67. Shapiro DD, Virumbrales-Munoz M, Beebe DJ, Abel EJ. Models of Renal Cell Carcinoma Used to Investigate Molecular Mechanisms and Develop New Therapeutics. Front Oncol. 2022;12:871252.

68. Liao TT, Yang MH. Hybrid Epithelial/Mesenchymal State in Cancer Metastasis: Clinical Significance and Regulatory Mechanisms. Cells. 2020;9(3).

69. Pastushenko I, Blanpain C. EMT Transition States during Tumor Progression and Metastasis. Trends Cell Biol. 2019;29(3):212–26.

70. Bostrom AK, Moller C, Nilsson E, Elfving P, Axelson H, Johansson ME. Sarcomatoid conversion of clear cell renal cell carcinoma in relation to epithelial-to-mesenchymal transition. Hum Pathol. 2012;43(5):708–19.

71. Ro JY, Ayala AG, Sella A, Samuels ML, Swanson DA. Sarcomatoid renal cell carcinoma: clinicopathologic. A study of 42 cases. Cancer. 1987;59(3):516-26.

72. He H, Magi-Galluzzi C. Epithelial-to-mesenchymal transition in renal neoplasms. Adv Anat Pathol. 2014;21(3):174–80.

73. Li R, Ferdinand JR, Loudon KW, Bowyer GS, Laidlaw S, Muyas F, et al. Mapping single-cell transcriptomes in the intra-tumoral and associated territories of kidney cancer. Cancer Cell. 2022;40(12):1583–99 e10.

74. Braun DA, Street K, Burke KP, Cookmeyer DL, Denize T, Pedersen CB, et al. Progressive immune dysfunction with advancing disease stage in renal cell carcinoma. Cancer Cell. 2021;39(5):632–48 e8.

75. Chakiryan NH, Kimmel GJ, Kim Y, Hajiran A, Aydin AM, Zemp L, et al. Spatial clustering of CD68+ tumor associated macrophages with tumor cells is associated with worse overall survival in metastatic clear cell renal cell carcinoma. PLoS One. 2021;16(4):e0245415.

76. Soupir AC, Hayes MT, Peak TC, Ospina O, Chakiryan NH, Berglund AE, et al. Increased spatial coupling of integrin and collagen IV in the immunoresistant clear cell renal cell carcinoma tumor microenvironment. bioRxiv. 2023.

77. Stringer C, Wang T, Michaelos M, Pachitariu M. Cellpose: a generalist algorithm for cellular segmentation. Nat Methods. 2021;18(1):100–6.

78. Pachitariu M, Stringer C. Cellpose 2.0: how to train your own model. Nat Methods. 2022;19(12):1634–41.

79. Danaher P, Zhao E, Yang Z, Ross D, Gregory M, Reitz Z, et al. Insitutype: likelihood-based cell typing for single cell spatial transcriptomics. bioRxiv. 2022:2022.10.19.512902.

80. Hansen J, Sealfon R, Menon R, Eadon MT, Lake BB, Steck B, et al. A reference tissue atlas for the human kidney. Sci Adv. 2022;8(23):eabn4965.

81. Vasconcelos AG, McGuire D, Simon N, Danaher P, Shojaie A. Differential Expression Analysis for Spatially Correlated Data. bioRxiv. 2024:2024.08.02.606405.

82. Yu G. Thirteen years of clusterProfiler. Innovation (Camb). 2024;5(6):100722.

83. McGue JJ, Edwards JJ, Griffith BD, Frankel TL. Multiplex Fluorescent Immunohistochemistry for Preservation of Tumor Microenvironment Architecture and Spatial Relationship of Cells in Tumor Tissues. Methods Mol Biol. 2023;2660:235–46.

84. Masotti M, Osher N, Eliason J, Rao A, Baladandayuthapani V. DIMPLE: An R package to quantify, visualize, and model spatial cellular interactions from multiplex imaging with distance matrices. Patterns (N Y). 2023;4(12):100879.

85. Foroutan M, Bhuva DD, Lyu R, Horan K, Cursons J, Davis MJ. Single sample scoring of molecular phenotypes. BMC Bioinformatics. 2018;19(1):404.

86. Chen Y, Chen L, Lun ATL, Baldoni PL, Smyth GK. edgeR v4: powerful differential analysis of sequencing data with expanded functionality and improved support for small counts and larger datasets. Nucleic Acids Res. 2025;53(2).

87. Therneau TM, Grambsch PM. Modeling Survival Data: Extending the Cox Model. 1st edition 2000. ed. New York, NY: Springer New York : Imprint: Springer; 2000. 1 online resource (XIV, 350 pages) p.

88. Kassambara A, Kosinski M, Biecek P. survminer: Drawing Survival Curves using ‘ggplot2’. 2024.

